# Oral Fumarate-based drugs alter gut microbiota species via cysteine succination

**DOI:** 10.64898/2026.01.08.698357

**Authors:** Marie Anselmet, Anna G. Burrichter, Thibaut Douché, Erica Bonazzi, Jean-Michel Betton, Bryan Jimenez-Araya, Mariette Matondo, Barbara Stecher-Letsch, Benoit Chassaing, Frédéric Barras, Rodrigo Arias-Cartin

## Abstract

Mono- and dimethyl fumarates are oral fumarate esters widely prescribed for relapsing-remitting multiple sclerosis and psoriasis. While these drugs appear to be effective, their effects on the gut microbiota and their precise bacterial targets remain unclear. In this study, we investigated how these drugs affect bacteria through a chemical modification called succination, where they react with protein thiol groups (-SH), particularly in cysteine residues. Using proteomics, enzymology and microscopy, we show how this post-translational modification disrupts several key bacterial functions and triggers oxidative and protein stress in *E.coli*. We also found that fumarate esters can be toxic to various gut bacteria in isolated cultures. Notably, our results demonstrate that when bacteria are studied together in microbial communities, the effect of fumarates can change, either weakening or intensifying. Our findings thus shed light on fumarate esters and microbiota interactions and allow identifying new molecular targets of succination relevant to microbiome health.

## INTRODUCTION

Post-translational modification (PTM) refers to the chemical and covalent change of amino acid side chains within proteins that alter their activity, structure, localization, and interactions. Hundreds of different PTMs exist, either enzymatically catalyzed or arising spontaneously from reaction with metabolites and xenobiotics^1^. Among these PTMs, cysteine succination has draw particular interest^2^ for its links to mitochondrial dysfunction and metabolic diseases such as hereditary leiomyomatosis and renal cancer^3^.

Fumarate esters (FAEs) such as Dimethyl Fumarate (DMF) and Monomethyl Fumarate (MMF) are used as intestinal-released medications against relapsing-remitting multiple sclerosis and psoriasis^4^. DMF and MMF either penetrate directly the intestinal epithelium or DMF is pre-metabolized into MMF by carboxylesterases before absorption^5^. Once incorporated FAEs inhibit signaling proteins such as NF-κB and block glycolysis and cell death^6–8^. At the biochemical level, MMF and DMF modify proteins by converting cysteine (Cys) residues into succinated derivatives, S-(2-methylsuccino)-cysteine (MSC) and S-(2-dimethylsuccino)-cysteine (DSC), respectively^5^. Succination was first described as the non-enzymatic reaction of fumarate with thiol groups *via* a Michael addition reaction, primarily those of Cys residues and free L-cysteines, leading to the formation of 2-(Succino)cysteine (2SC) adducts^9^ (Figure 1A). It was described in conditions where fumarate - a metabolite of the tricarboxylic acid (TCA) cycle - levels rise, such as in Fumarate hydratase-deficient cells^10^. Interestingly, the formation of 2SC being irreversible under physiological conditions, 2SC serves as a biomarker for dysfunctional physiological states and several diseases^11^. Similarly, succination by FAEs has gathered significant attention for its immunomodulatory and anti-inflammatory potential. Beyond its immunosuppressive properties, emerging research suggests that DMF could also exert effects on the intestinal microbial community. However, FAEs-microbiota interactions, as well as the exact mechanism at play remain unclear, frequently leading to contradictory conclusions across studies and yielding results with low reproducibility^12–14^.

**Figure 1.**
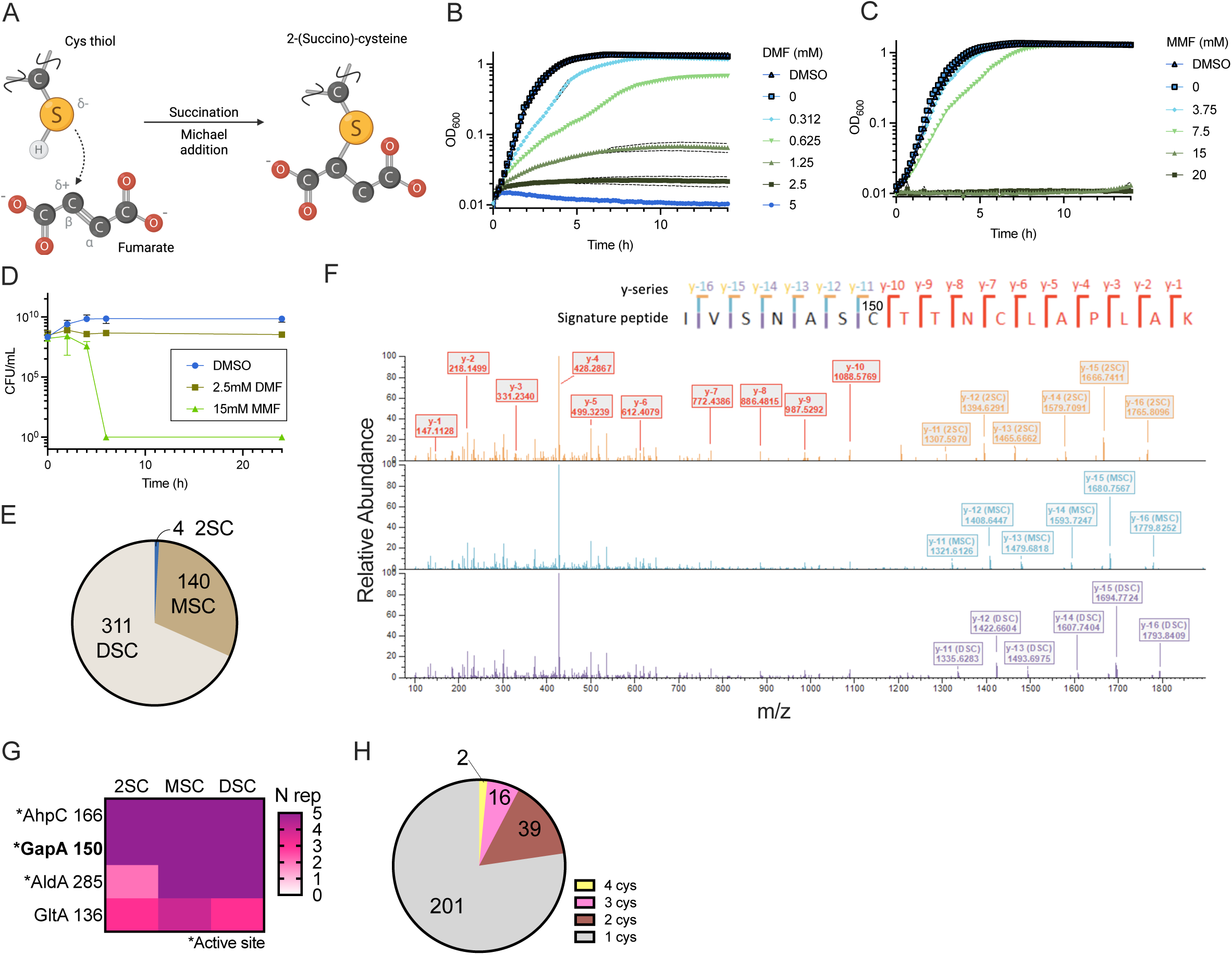
DMF and MMF intoxicate *E. coli* through succination. (A) Succination of cysteine thiol by Michael addition reaction with fumarate. (B and C) Growth of *E. coli* exposed to DMF (A) and MMF (B) in LB medium aerobically. DMSO control was carried out at 4% v/v, equivalent to the highest FAE concentration tested. n = 2 replicates from 3 biological replicates. Standard deviation is represented in black dotted line. (D) Colony Forming Unit (CFU) assay on *E. coli* after exposure to 2.5 mM DMF or 15 mM MMF. DMSO = 3% v/v. n = 3 technical replicates from 3 biological replicates per treatment. Standard deviation is represented with bars. (E) Distribution of succination species identified in the proteome of *E. coli* exposed to DMF. (F) MS2 fragmentation spectra of the GapA signature peptide YAGQDIVSNASC_150_TTNCLAPLAK peptide according to the different types of succination: 2SC (top spectrum), MSC (middle spectrum) and DSC (bottom spectrum) species. The three spectra contain 2 fragments: -TTNCLAPLAK of the peptide with red inserts/rectangles, which represent the non-modified and equivalent fragmentation pattern, and YAGQDIVSNASC_150_- with orange, turquoise and purple inserts, which are directly linked to the mass of the succinations. (G) Succination species in AhpC Cys_166_, GapA Cys_150_, AldA Cys_285_ and GltA Cys_136_. Color indicates the number of replicates were the PTM was detected out of 5 biological replicates. (H) Distribution of number of Cys^suc^ per protein.

Here we report the use of *Escherichia coli*, the Oligo-Mouse-Microbiota (OMM^12^) model as well as *in vitro* human gut microbiota communities modelization to study the interplay between DMF/MMF and members of the intestinal microbiota. First, we investigated by mass spectrometry-based proteomics the effect of DMF in *E. coli*. Our approach highlighted the PTM of hundreds of Cys residues into MSC and DSC adducts in the *E. coli* proteome that led to the inactivation of essential metabolic enzymes. Furthermore, we uncovered the *E. coli* proteomic and physiological response during DMF exposure using *in vivo* and *in vitro* tests. Finally, we assessed the effect of DMF and MMF on the composition of two *in vitro* model microbiota communities. Our study reports that those drugs are toxic for several microbiota members in isolated cultures, and that such toxic effects can be damped or exacerbated in consortia. Overall, this study provides new insights into the targets of highly prescribed fumarate esters, their mode of action *in vivo* and identifies physiological consequences on gut bacterial inhabitants.

## RESULTS

### Physiological effect of DMF and MMF treatments in E. coli

To study the effects of FAEs on bacteria, we exposed *E. coli* MG1655 to DMF and MMF in LB liquid medium and performed growth curves as well as viability plate assays aerobically. In liquid cultures we observed adverse effects on *E. coli* growth starting at 0.312 mM DMF and 7.5 mM MMF (Figure 1B and 1C, respectively). Total growth inhibition was observed at 2.5 mM DMF (Figure 1B) and 15 mM MMF (Figure 1C). Interestingly, through viability assays (Figure 1D) using 2.5 mM DMF or 15 mM MMF, we showed a low killing effect of DMF and a full bactericidal effect of MMF after 6h of exposure. Dimethyl Sulfoxide (DMSO) control used for DMF and MMF solubilization showed no impact on growth. Together, these results showed that both FAEs have an antibacterial effect, DMF being bacteriostatic and MMF bactericidal.

### Detecting succinated proteins in DMF-treated E. coli

We next performed a proteomic analysis of *E. coli* cultures exposed to 2.5 mM DMF during 2h. We detected 455 succinated peptides across three forms: 140 MSC, 311 DSC and only 4 2SC (Figure 1E and Table S1). The 2SC succinated peptides were Cys_166_ of the Alkyl hydroperoxide reductase (AhpC), Cys_150_ of the Glyceraldehyde 3-phosphate dehydrogenase A (GapA or GADPH, Figure 1F), Cys_285_ of the Aldehyde dehydrogenase A (AldA) and Cys_136_ of the Citrate synthase (GltA) (Figure 1G). According to GOTerm enrichment analysis (Table S2 and Figure S1), the 258 succinated proteins were part of many diverse pathways and many had catalytic function or were cofactor dependent. Strikingly, many proteins belonging to metabolic pathways such as glycolysis and TCA cycle were succinated (Table S1). In particular, GapA, an essential enzyme for *E. coli* involved in glycolysis^15^ had 3 succinated Cys residues (Cys^suc^) (Cys_150_, Cys_154_ and Cys_289_). Of note, proteins with the highest number of identified Cys^suc^ were GltA and the tryptophanase / L-cysteine desulfhydrase (TnaA) with 4 Cys^suc^ each, while 16 other proteins had 3 Cys^suc^ (Figure 1H and Table S1). Thus, FAE cause irreversible succination of Cys residues involved in catalytic activity or binding cofactors, and across a wide range of structurally and functionally diverse enzyme types. Moreover, it demonstrates that DMF is pre-metabolized into MMF in bacteria.

### Assessing changes in steady-state level of cellular proteins in DMF-treated E. coli

To determine the effects of DMF on *E. coli* we conducted a comparative proteomic analysis of cells treated with DMF or DMSO. A total of 127 proteins showed enhanced level or were exclusively detected in the DMF-treated sample while 309 proteins were less abundant or absent (Figure 2A and Table S3). GOTerm enrichment analysis comparing DMF vs DMSO treated cells indicated that abundance of [Fe-S] cluster containing proteins -AcnB, FumA, FumB, IspH and PreT- (Figures 2B and 2C), of metal bound proteins (Figure 2C), and of proteins involved in respiration or energy production were absent or in lower abundance in the DMF-treated sample (Table S2). On the opposite, DMF treatment enhanced the level of proteins involved in oxidative stress adaptation such as AhpC, AhpF, GrxA, KatE and TrxC and response to heat stress such as DnaK, ClpB, IbpA, IbpB, HslU and HslV (Figure 2B and 2D). Last, DMF treatment enhanced levels of proteins involved in the in L-Arginine degradation, sulfur homeostasis, and stress adaptation responses, including IscR and MarR transcriptional regulators (Figures 2B, 2D and 2E).

**Figure 2.**
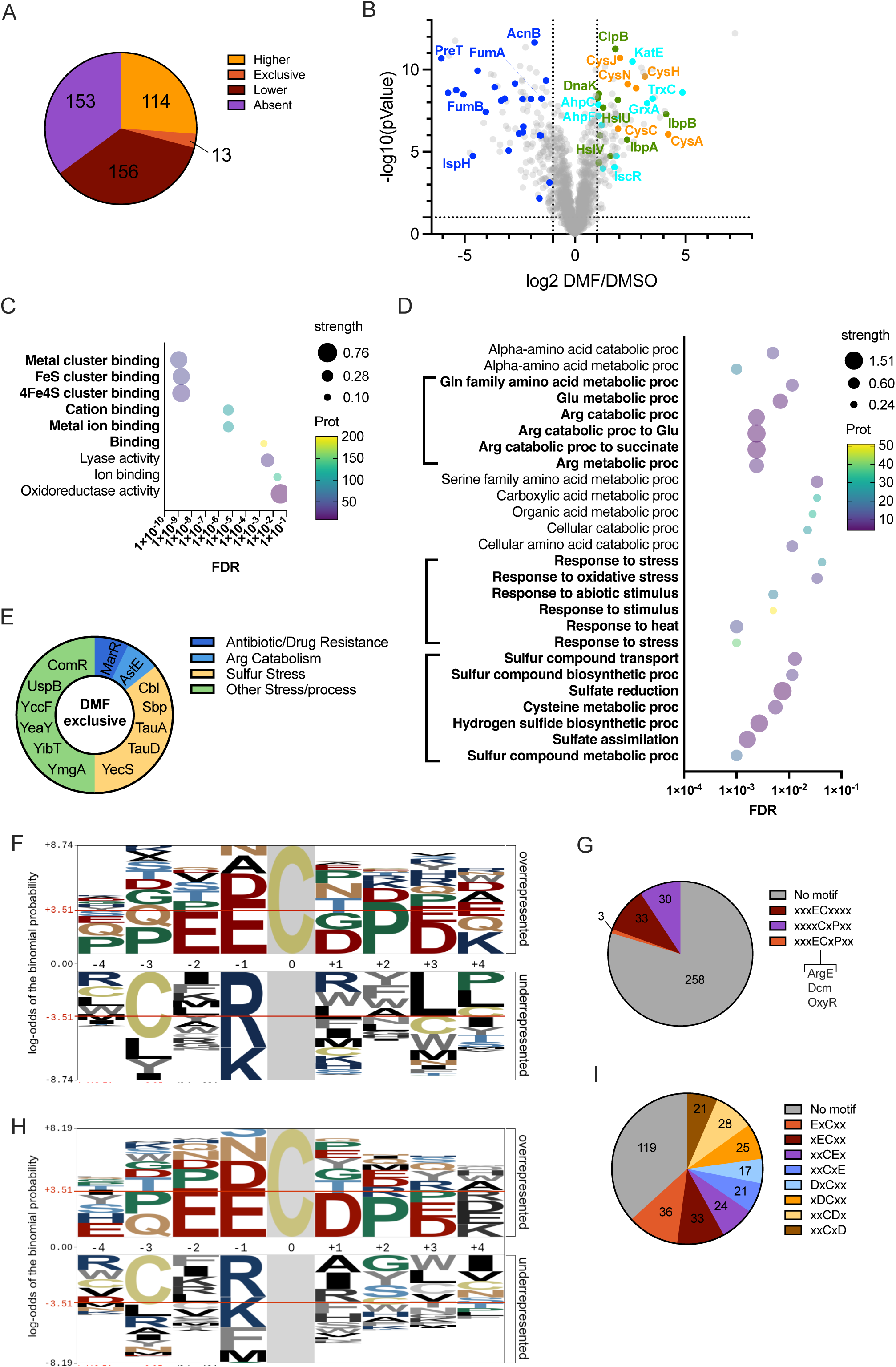
*E. coli* proteomic response towards DMF. (A) Differential abundance analysis of proteins identified in *E.coli* DMF-exposed vs DMSO control. Proteins absent, exclusive or with higher/lower abundance with absolute log_2_ (fold-change) superior to |1| in the DMF-treated sample. (B) Differentially expressed proteins of the folding stress response (green), redox (cyan) and sulfate assimilation pathways (orange) or [Fe-S] cluster proteins (blue). Dotted lines indicate log_2_ fold =1 and -log_10_ (pValue) = 1 threshold. (C) GOTerm molecular function enrichment analysis of proteins absent or with lower abundance in DMF-treated cells.* (D) GOTerm process function enrichment analysis of proteins with higher abundance or unique in DMF-exposed *E. coli* cells.* (E) GOTerm process function categories of proteins exclusively detected in the DMF-exposed *E. coli* - sample. (F) pLOGO motif analysis on MSC and DSC Cys^suc^ from DMF-treated *E. coli*. (G) Distribution of ECxP motif in the MSC and DSC Cys^suc^ peptides. (H) pLOGO motif analysis on DSC Cys^suc^ from DMF-treated *E. coli*. (I) Distribution of negative charged residues at -2 or +2 residues from the DSC Cys^suc^ peptides. * FDR: False Discovery Rate (p-values corrected for multiple testing within each category using the Benjamini–Hochberg procedure). Prot scale bar: color indicates the number of proteins within a particular category. Strength: Log_10_(observed /expected).

### Towards defining a Cysteine Succination pattern

Previous studies have searched for signatures or patterns that could predict Cys-containing motifs that are likely targets of fumarate, MMF or DMF^16,17^. We investigated if such information could be extracted from the analysis of linear sequences flanking Cys^suc^ containing proteins. By using the pLOGO statistical tool^18^ we scanned a 4 residues window encompassing the 324 Cys^suc^ identified. This motif analysis showed enrichment of glutamate (E) or proline (P) near the Cys^suc^ residues, forming an ECxP motif (Figure 2F), which occurred in only 3 peptides (from ArgE, Dcm and OxyR proteins), and therefore no predictive (Figure 2G). Still almost 10% of peptides were identified with an ECxx motif or a xCxP motif (Figure 2G). When the pLOGO analysis was obtained from peptides that contained only DSC, a second E residue was found associated with the Cys^suc^ (EECxx, Figure 2H). Overall, 205 Cys^suc^ have at least one negative charged residue (D or E) within +2 and -2 residues from the Cys^suc^ (Figure 2I). Altogether, this analysis revealed a primary sequence trend identifying potentially succinated sites, yet it failed to identify a highly discriminant succination consensus sequence.

### DMF/MMF effects on GADPH and selected TCA enzymes

Next, we used biochemical assays to further investigate the effects of FAEs exposure on specific enzymes pointed out in the proteomic analysis. Cell cultures were exposed to 2.5 mM DMF or 15 mM MMF for 2h and cell extracts were prepared. The activity of glycolysis enzyme GADPH was almost completely abolished (Figure 3A) and activities of the TCA enzymes aconitase and fumarase were also impaired or greatly decreased (Figures 3B and 3C). The MS-based analysis revealed that Cys^suc^ in GAPDH was the catalytic Cys_150_ residue while Cys^suc^ in aconitase and fumarase were Cys residues acting as ligands for [Fe-S] cluster (Table S1 and Figures 3A, 3B and 3C). Interestingly, DMF/MMF treated extracts showed no decrease in succinate dehydrogenase activity (Figure 3D). Consistently, only the surface-exposed SdhA Cys_303_ was succinated in this enzymatic complex (Table S1 and Figure 3D), which has no known involvement in its activity. These results provided a molecular rationale of the inactivation of [Fe-S] cluster bound enzymes targeted by succination.

**Figure 3.**
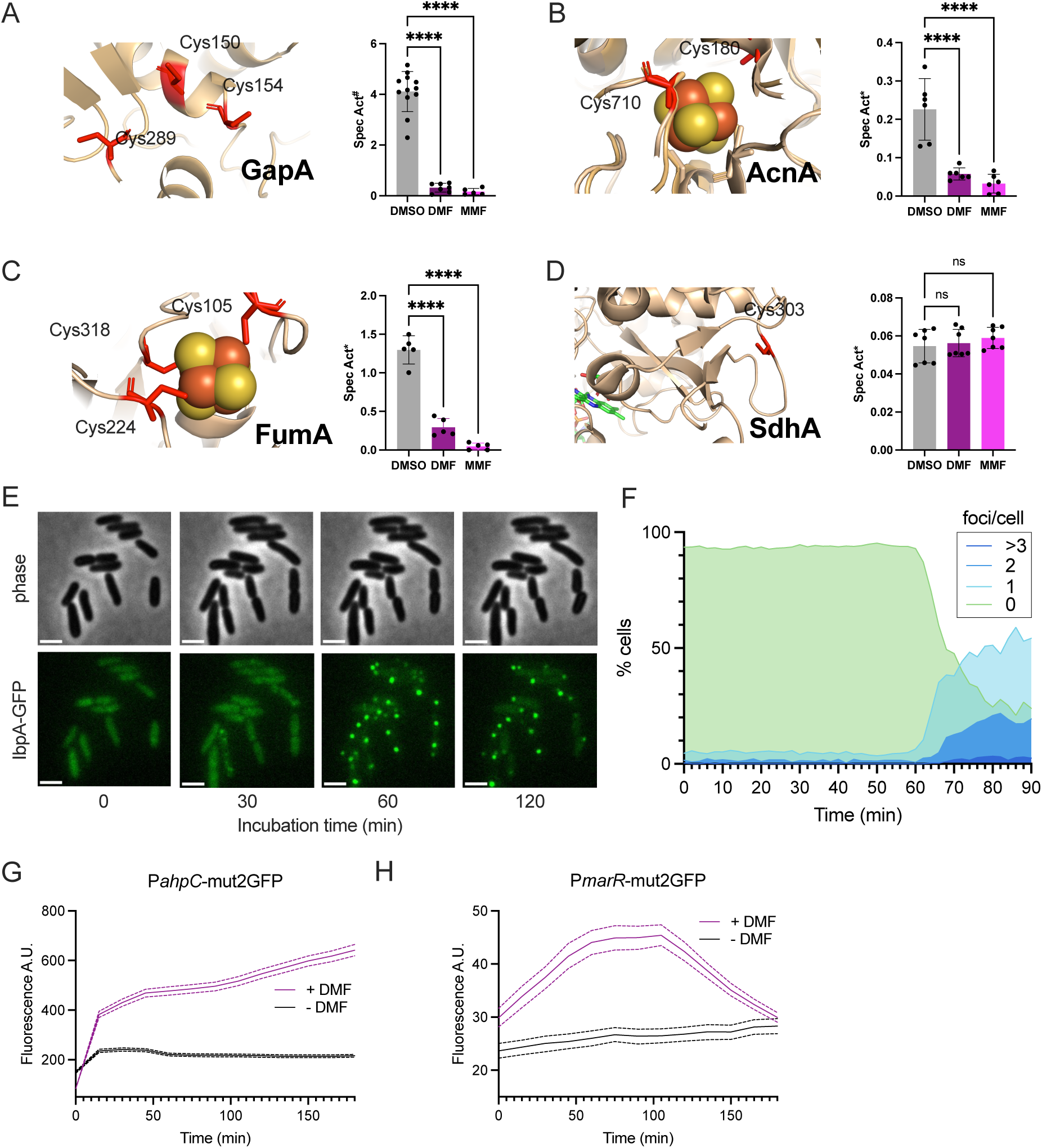
Physiological effects of succination in glycolysis and TCA cycle. (A-D) Enzymatic specific activity of GAPDH (A), aconitase (B), fumarase (C) and succinated dehydrogenase (D) in *E. coli* cell lysates exposed to FAEs. Left panels depict the tridimensional structures for GAPDH (pdb:1DC3), aconitase B (pdb:1L5J), fumarase A (AF-P0AC33-F1-v4 model) and succinate dehydrogenase (pdb:1NEK). Right panels indicate specific activity in mmol NADH (A), μmol aconitate (B), μmol fumarate, (C) and nmol NADPH, (D) **⋅**min^-1^**⋅**mg^-1^. n = at least 5 replicates from 2 biological replicates. p values determined by one-way ANOVA analysis within the indicated pairs are indicated by asterisks: **p < 0.01, ***p < 0.001, ****p < 0.0001 and n. s. = not significant. (E) Microscopic time lapse images of *E. coli* cells expressing an IbpA-msfGFP fusion and exposed to 2.5 mM DMF in M9glycerol. Scale bar: 2μm (F) Analysis of time lapse experiment of cells producing IbpA-msfGFP foci upon DMF exposure. (G) and (H) analysis of time lapse experiment of cells expressing P*ahpC*-mut3GFP or P*marR*-mut3GFP in presence or absence of 2.5 mM DMF.

### DMF causes proteotoxicity

The above proteomic analysis revealed that inclusion body-associated protein A and B -IbpA and IbpB-were more abundant in the DMF-exposed cells (Figure 2B and Table S3). Those proteins belong to a conserved class of α-crystallin ATP-independent chaperones that bind to unfolded protein substrates and prevent their irreversible aggregation in stress conditions (e.g. heat shock, oxidative stress or antibiotics)^19^. Hence, we suspected that DMF-treated cells underwent a stress-induced unfolding or misfolding of succinated proteins and/or defects in protein translation. In support of this view, we detected Cys^suc^ in ribosomal subunits (RplJ, RpmA, RpsL), in proteins involved in ribosome biogenesis (RimO, RmsL and RmsB) and t-RNA modification (MmnA, MnmG, ThiI, TsaB and YgfZ) (Table S1). Therefore, to investigate the effect of DMF on protein folding, we designed an *ibpA*-msfGFP fusion at gene locus in *E. coli* and followed its response to DMF by microscopy. In M9 glycerol medium supplemented with 2.5 mM DMF we observed the IbpA-msfGFP fusion homogenously distributed in the cytoplasm (Figure 3E) at t = 0 min. Then, starting at 50-60 min of DMF exposure cells formed intracellular foci (Figures 3E and 3F), which pointed out protein folding stress^20^. Most of the cells formed one focus, although 2 or >3 foci per cell could be observed. Cells with at least one focus reached up to 80% of the population at 90 min. In the control experiment with DMSO only <0.8% of cells had GFP aggregates (Figure S2). Thus, these single cell microscopy-based experiments showed that DMF triggers protein folding stress in *E. coli*.

### DMF enhances oxidative stress

Enhanced level of anti-ROS proteins (Figure 2B, AhpC, AhpF, GrxA, IscR, KatE and TrxC) suggested that DMF treatment induces an oxidative stress response. Hence, we performed time lapse microscopy experiments on cells harboring fluorescent transcriptional reporters for *ahpC* gene expression, which is under control of the oxidative stress sensing regulator OxyR (<u>Ox</u>idative stress <u>R</u>egulator)^21^. Consistent with an intracellular ROS production, the expression of both *PahpC*-mut2GFP increased upon DMF exposure (Figure 3G). Remarkably, the fluorescence from *PahpC*-mut2GFP quickly increased in the first 30 min of the experiment, meaning that DMF may produce ROS rapidly. Furthermore, we tested the expression of the *marR* gene, which is under control of SoxRS (<u>S</u>uper<u>ox</u>ide Response proteins) -another major oxidative stress regulator. We observed that, fluorescence produced from a *PmarR*-mut2GFP fusion rose over 100min in the presence of DMF and later decreased for reasons that remain unclear (Figure 3H). These results showed that DMF treatment produces enhanced levels of ROS.

### DMF targets sulfur metabolism

Proteomic analysis revealed that exposure to DMF increased the abundance of proteins of the sulfur assimilation/L-cysteine biosynthesis pathways (Cys,Tau and Ssu) in *E. coli* (Figures 2B, 2E and Table S2). We hypothesized that this response was derived from exhaustion of L-cysteine upon succination by DMF/MMF forming 2SC adducts. Therefore, we tested whether addition of sulfur sources, L-cysteine or taurine, to the medium could abrogate DMF toxicity. Indeed, we observed that addition of L-Cys at 0.5mM or 0.05mM restored cell growth (Figure 4A, top panel). However, taurine did not rescue from DMF toxicity at any of the concentrations used (Figure 4A, bottom panel). L-cysteine is the source of glutathione (GSH) (Figure 4B). Therefore, we quantitated the level of reduced and oxidized glutathione (GSH/GSSG) in DMF exposed cells and observed that DMF treated cells exhibited a strong GSH/GSSG depletion (Figure 4C). Last, PTMs analysis by MS showed that GshA, which catalyzes GSH formation, had its catalytic Cys residue succinated after DMF treatment (Figure 4B and Table S1), providing an additional rationale for GSH depletion. Overall, this set of experiments highlighted that DMF exposure altered Cys, redox and sulfur homeostasis. It is interesting to note that DMF-exposed cells synthesized a lower percentage of Cys containing proteins as compared with untreated cells (Figure 4D). Presumably, like observed in sulfur deficient cells^22^, optimized allocation of Cys and/or sulfur resources is taking place under DMF stress.

**Figure 4.**
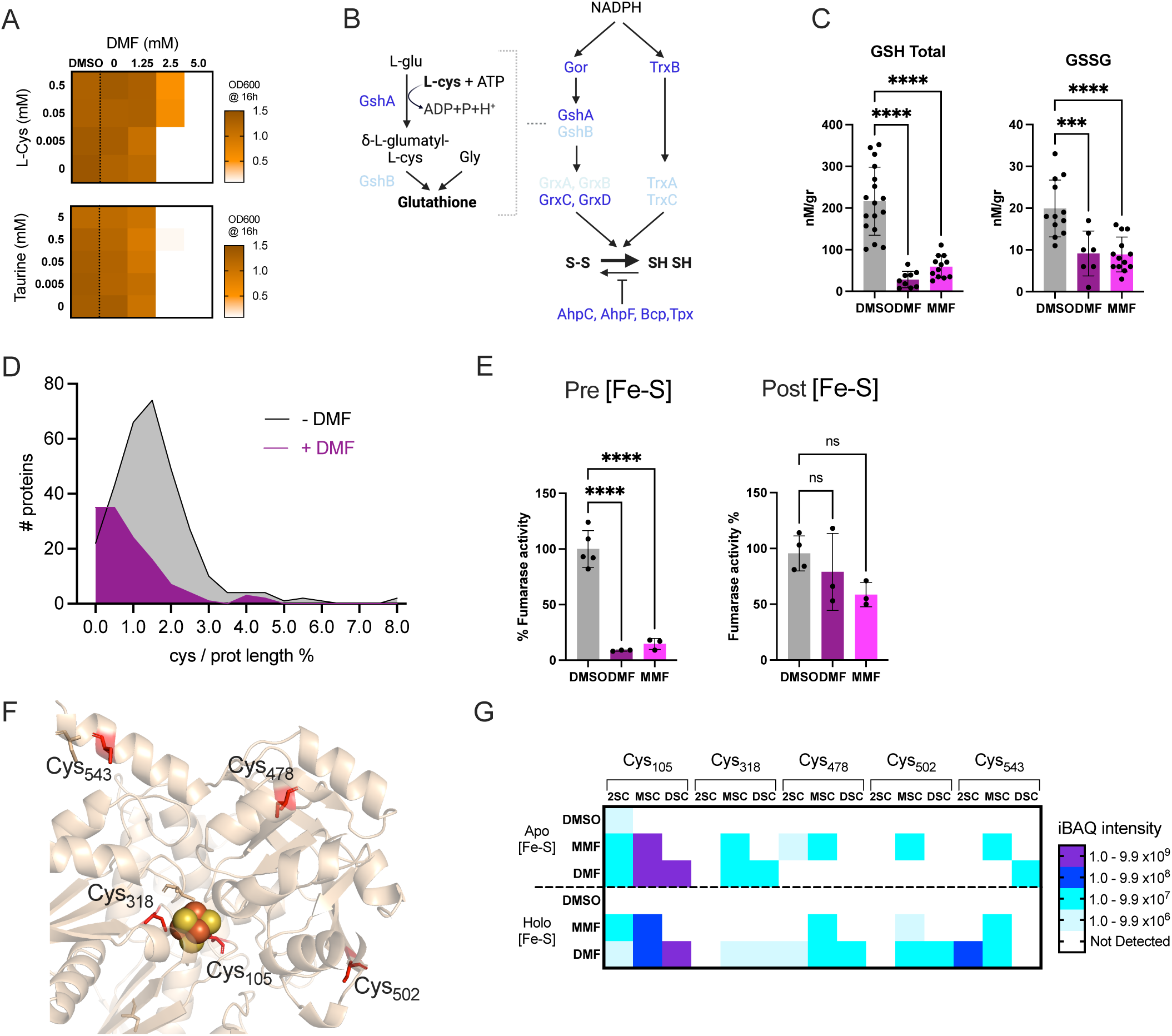
DMF induces oxidative stress, targets sulfur metabolism and apo [Fe-S] proteins. (A) Effect of L-cysteine (top panel) and Taurine (bottom panel) supplementation on cells exposed to DMF. (B) Glutathione and redox homeostasis pathways in *E. coli*. Arrows: Reaction direction. Purple: Succinated proteins. Light blue: No succinated proteins. Black: Reaction substrates and products. (C) Reduced (GSH) and oxidized glutathione (GSSG) in *E. coli* cells exposed to DMF, MMF or DMSO control. n = at least 7 replicates from 2 biological replicates. Standard deviation is represented with bars p values determined by one-way ANOVA analysis within the indicated pairs are indicated by asterisks: **p < 0.01, ***p < 0.001, ****p < 0.0001 and n. s. = not significant. (D) Distribution of number of Cys^suc^ per protein. (E) Enzymatic activity of purified FumA exposed to FAEs before (Pre [Fe-S] ) or after (Post [Fe-S] ) [Fe-S] cluster *in vitro* reconstitution. Activity normalized to the mean of DMSO control. n = at least 3 replicates. p values determined by one-way ANOVA analysis within the indicated pairs are indicated by asterisks: **p < 0.01, ***p < 0.001, ****p < 0.0001 and n. s. = not significant. (F) Position of Cys^suc^ (red sticks) in FumA tridimensional model (AF-P0AC33-F1-v4 model). (G) Cys^suc^ detected by LC MS/MS in apo and holo FumA exposed to DMSO (control), MMF, or DMF.

### DMF targets [Fe-S] clusters proteins

Proteomic profiling showed that [Fe-S] cluster-bound proteins have a much lower abundance in the DMF-treated cells (Figures 2B and 2C, and Tables S3 and S4). We also noticed that several components involved [Fe-S] biogenesis, i.e. IscU, Fdx, IscA and GrxD were found to be succinated (Table 1). Noticeably, the cluster of most of the succinated [Fe-S] proteins located in solvent exposed positions. Therefore, we hypothesized that lower abundance of those [Fe-S] proteins resulted from succination of Cys residues serving as [Fe-S] cluster ligands, which would preclude [Fe-S] cluster insertion and eventual degradation of the apo-form (without its [Fe-S] cluster). In support of this, we observed that twelve [Fe-S] cluster proteins showed Cys^suc^ involved in [Fe-S] coordination (Table 1). Notably, FumA was succinated at all its [Fe-S] cluster coordinating Cys residues (Table 1 and Figure 3C). To test whether succination could prevent [Fe-S] insertion, we next purified the FumA apo-form and followed its activity before or after the reconstitution of its [Fe-S] cluster (holo-form) in the presence or absence of DMF or MMF. Again, for these assays, we used a 1:9 molar ratio of FumA to DMF or MMF, since FumA has 9 Cys^suc^. When the Apo-FumA was preincubated with FAEs before the [Fe-S] cluster reconstitution, only a small fraction of the resulting protein was active (Figure 4E, Pre [Fe-S]). At the opposite, 2/3 of protein activity remained when the holo-FumA was incubated with FAEs in the same conditions (Figure 4E, Post [Fe-S]). The LC-MS/MS analyses of trypsin-digested FumA derived from these *in vitro* experiments detected 5 Cys^suc^ residues on this sample, of which 2 are involved in [Fe-S] coordination (Figure 4F, Cys_105_ and Cys_318_) and the rest are predicted to be solvent-exposed residues (Figure 4F, Cys_478_, Cys_502_ and Cys_543_). In general, succination of Cys_105_ and Cys_318_ in the FAE-exposed samples is not detected or with a lower peptide intensity on the holo-FumA compared to the apo-FumA (Figure 4G and Table S4). Therefore, DMF can compete with [Fe-S] insertion through succination of the liganding Cys residues at solvent-exposed [Fe-S] cluster binding sites. However, our experiments also showed that bound [Fe-S] clusters protect Cys residues ligands from succination by DMF and MMF.

**Table 1.** Succinated FeS biogenesis machinery factors and [Fe-S] cluster proteins.

| [Fe-S]<br>protein<sup>a</sup> | Succination<sup>b</sup> |  |  | Cys function | [Fe-S] solvent<br>exposed |
| --- | --- | --- | --- | --- | --- |
|  | Cys | MSC | DSC |  |  |
| AcnA | 180 | 2 | 3 | Unknown | Yes |
| AcnB | 710 | 0 | 2 | [Fe-S] ligand | Yes |
| Fdx<sup>c</sup> | 13 | 0 | 2 | [Fe-S] ligand | Yes |
| FumA | 105 | 3 | 5 | [Fe-S] ligand | Yes |
| FumA | 224 | 0 | 5 | [Fe-S] ligand | Yes |
| FumA | 318 | 1 | 2 | [Fe-S] ligand | Yes |
| GrxD<sup>c</sup> | 30 | 0 | 3 | [Fe-S] ligand | Yes |
| IlvD | 395 | 2 | 0 | Unknown |  |
| IscA<sup>c</sup> | 101 | 0 | 3 | [Fe-S] ligand | Yes |
| IscA<sup>c</sup> | 99 | 0 | 2 | [Fe-S] ligand | Yes |
| IscU<sup>c</sup> | 37 | 1 | 0 | [Fe-S] ligand | Yes |
| MnmA | 102 | 4 | 4 | [Fe-S] ligand | Yes |
| PflA | 13 | 0 | 2 | [Fe-S] ligand | Yes |
| RimO | 17 | 0 | 3 | [Fe-S] ligand | Yes |
| RlmD | 389 | 0 | 3 | unknown |  |

<sup>a</sup>[Fe-S] proteins according to Lenon et al<sup>33</sup>
<sup>b</sup>Number of samples detected were succination was detected within 5 biological replicates.
<sup>c</sup>Proteins of the ISC [Fe-S] biogenesis machinery according to Py et al<sup>34</sup>

### DMF and MMF intoxicate select mouse intestinal microbiota bacterial members

DMF and MMF are active ingredients of orally administered drugs with delayed release within the gastrointestinal tract. Therefore, we next aimed to extend our analysis of FAEs effects to bacterial species that are likely to be found in the gut. For this purpose, we first employed the Oligo Mouse Microbiota community (OMM^12^), a synthetic microbial community that consists of twelve characterized members of the mouse gut microbiome^23^. We tested the effect of DMF and MMF anaerobically on each of these 12 bacteria, as well as on *E. coli* MG1655, in individual cultures using AF medium^23^. Of note, MMF addition lowered slightly the pH of the AF medium. Therefore, as pH could alter growth of some OMM^12^ strains^24^ we readjusted medium pH to 7.0 when MMF was used.

We interestingly identified four main phenotypes. The first included species that were sensitive to DMF and MMF, and comprised *E. coli* (Eco), *Akkermansia muciniphila* (YL44), *Bacteroides caecimuris* (I48), *Muribaculum intestinale* (YL27) and *Turicimonas muris* (YL45) (Figures S3A, S3B, S3C, S3D and S3E respectively). The second class included species resistant to both FAEs, and comprised *Bifidobacterium animalis* (YL2), *Enterococcus faecalis* (KB1) and *Limosilactobacillus reuteri* (I49) (Figure S3F, S3G, and S3H, respectively). The third class included species that were resistant to DMF but sensitive to MMF, and comprised *Clostridium innocuum* (I46), *Enterocloster clostridioformis* (YL32) and *Flavinifractor plautii* (YL31) (Figures S3I, S3J and S3K, respectively). The fourth class includes *Blautia coccoides* (YL58) that showed full resistance against both DMF and MMF after several hours of exposure (Figures S3L). Besides, *Acutalibacter muris* (KB18) was found to be sensitive to DMSO, making assessment of its FAEs’s sensitivity impossible (Figure S3M). Last, the addition of DMSO stimulated *E. coli* growth (Figure S3A), likely due its capacity to metabolize this chemical via anaerobic respiration^25^. These results revealed that when treated individually, bacterial species of the mouse gut exhibit differential sensitivity to toxicity of FAEs.

Next, we aimed to investigate the toxicity of FAEs at the ecosystem level in the OMM^12^ consortium. Communities were inoculated with individual species mixed in equal proportions and then stabilized for five days with transfers into fresh medium every 24h. At this, point (D0) the community was exposed to FAEs for 3 additional days with a daily transfer into fresh medium. After 3 days (D1-D3), we performed qPCR analysis to determine absolute and relative abundance of the different OMM^12^ species (Figure 5A and Table S5). Absolute qPCR analysis on OMM^12^ consortium showed that DMF and MMF exerted a slight toxic effect on the OMM^12^ consortium (Figure S4A, total panel). Also, this qPCR analysis failed to identify YL2, KB18 and I49, presumably due to their loss during the equilibration step (Figure S4A). Exposure to DMF or MMF produced a high relative abundance of YL58 and a lower abundance of all the other strains (Figures 5B). Thus, tolerance of YL58 was equivalent to what we had observed in FAEs-treated isolated cultures (Figures S4L). In contrast, we observed that YL44, I48, YL27 and YL45 were able to tolerate FAEs when present in the OMM^12^ consortium (Figures 5B and S4A). Thus, these results suggest that OMM^12^ consortium could provide protection for some of the strains killed by one or both FAEs in single cultures.

**Figure 5.**
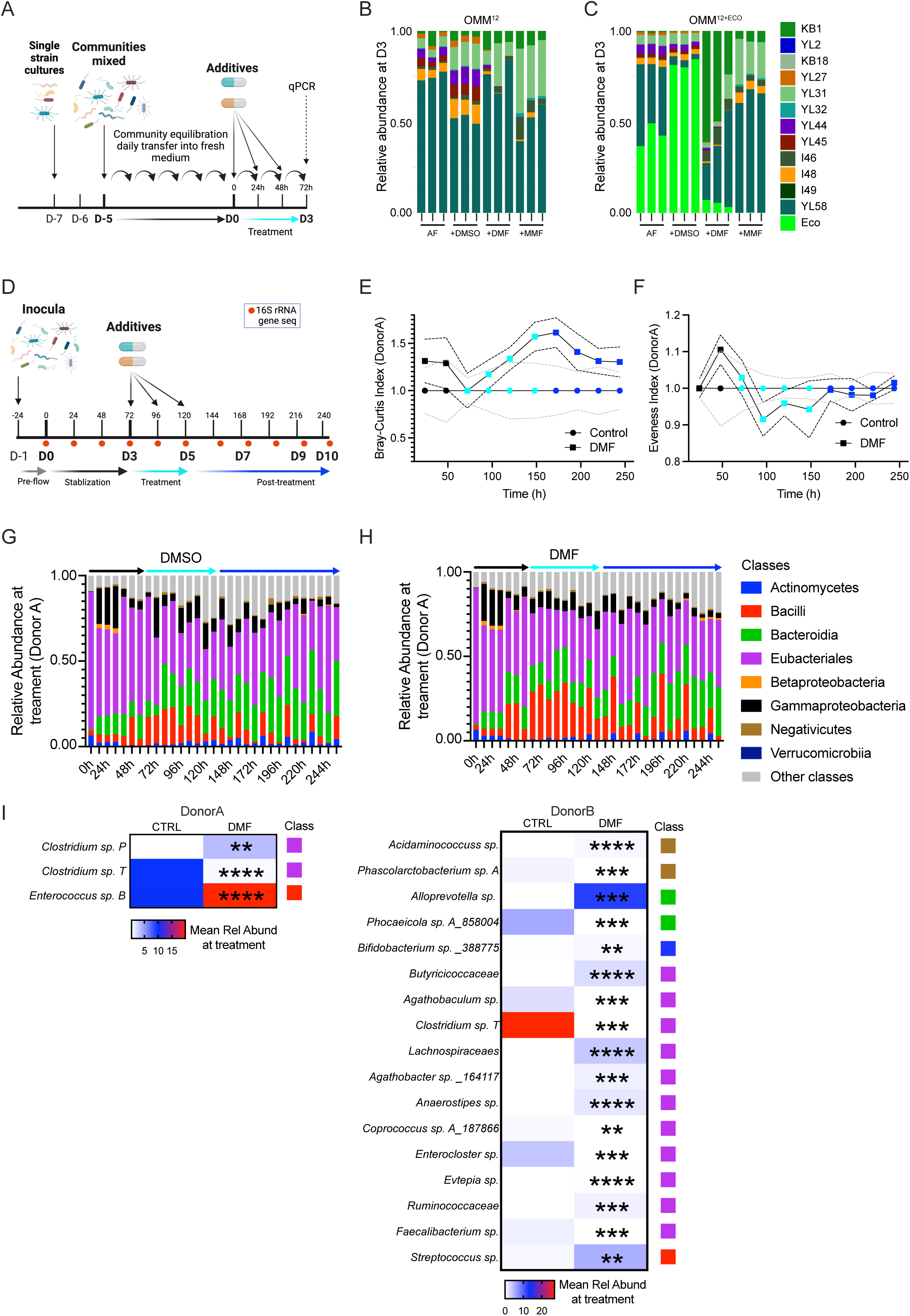
Dynamics of OMM and MBRA consortiums upon exposure to FAEs. (A) Scheme of the OMM^12^ and OMM^12+Eco^ community experiments. (B) Relative abundance of the OMM^12^ and (C) OMM^12+Eco^ microbiota consortiums during exposure after three days of treatment with DMF, MMF or controls. Abundance of each individual strain is shown as relative abundance and expressed as percentage of cumulative 16S rRNA gene copy numbers of all strains. One bar corresponds to one of three replicates.(D) Scheme of the MBRA community experiments. (E) Bray-Curtis distance and (F) Evenness index analysis of MBRA samples generated with the DMF and DMSO control from Donor A samples. Data were normalized to the DMSO control at each time point, defined as 1. Dots color indicates the stabilization, treatment and post-treatment points. Solid and dotted lines correspond to the mean and standard deviation of triplicates of each time point, respectively. (G) Relative abundance of the MBRA control and (H) DMF-exposed microbiota consortiums at Class taxonomic level during a 10-day long experiment. Abundance of each individual class is shown as relative abundance and expressed as percentage of cumulative 16S rRNA gene copy numbers of the rest of the classes. One bar corresponds to one of three replicates. Arrow color corresponds to the experiment section, stabilization (black), treatment (cyan) and posttreatment (blue). (I) Species with significantly different abundance between the DMSO control and DMF-treated samples during the 72h, 96h and 120h points for donor A (left panel) and donor B (right panel) . Square colors correspond to the classes which the species belongs. p values determined by two-way ANOVA analysis within the indicated pairs are indicated by asterisks: **p < 0.01, ***p < 0.001, ****p < 0.0001 and n. s. = not significant.

Since the OMM^12^ community seemed to provide fitness advantage in the presence of FAEs for a subset of species, we next tested whether the OMM^12^ community could also protect *E. coli* MG1655 against FAEs. This consortium, called hereafter OMM^12+Eco^, displayed absolute qPCR values reminiscent to the OMM^12^ upon exposure to FAEs, showing that DMF and MMF were toxic as well (Figure S4B). The qPCR-based abundance analysis revealed that *E. coli* represented 30-50% of the community in the AF control and up to 70-80% in the DMSO group (Figure 5C and S4B). Moreover, addition of DMF or MMF on this community produced a similar diversity to the OMM^12^ consortium (Figure 5B), except for *E. coli* that was also detected in the DMF sample (Figure 5C). As a reminder, *E. coli* was otherwise killed in isolated cultures (Figure S3A). Hence, this data importantly highlighted that the OMM^12^ community could offer protection to *E. coli* against DMF.

### DMF and MMF intoxicate human intestinal microbiota bacterial members

Collectively, the data above, pointed toward a toxic effect of FAEs compounds on several bacterial members of the mouse gastrointestinal tract, either in isolation or at the community level. Therefore, we next aimed to investigate the potential human relevance of these findings by using an *in vitro* simulator of the human intestinal microbiota. More specifically, we employed the MiniBioReactor Arrays (MBRA), an *in vitro* microbiota modelling system that allows dynamic and stable culture of human-derived microbiota under anaerobic conditions. Fecal samples from two healthy subjects were used to inoculate triplicate MBRA chambers and subjected to a stabilization, a treatment, and a post-treatment phase.

After a 3 days stabilization period, MBRA microbiotas were then treated daily for 3 days with DMSO as control or with DMF, followed by a 5 days-long post-treatment phase (Figure 5D). MBRA chambers were sampled daily over 10 days and assessed for microbiota composition *via* Illumina-based 16S rRNA gene sequencing. This approach importantly revealed temporal Bray-Curtis dissimilarity induced by DMF treatment, highlighting that DMF impacted the presence and/or relative abundance of various members of the modelized human intestinal microbiota (Figure 5E), with a non-significant effect on microbial alpha diversity, as revealed through evenness index computing (Figure 5F). Taxonomical analysis revealed that such DMF-induced alterations of the microbial ecosystem were impacting various members of the ecosystem, including a beneficial impact on the Bacilli class, such as *Enterococcus* species, as well as a negative impact on species from the *Clostridium* genus (Figure 5I left panel). Interestingly, a similar experimental approach was performed on an additional donor (Figure S5), revealing significant impact of DMF exposure on the obtained ecosystem but with impact on different microbiota members (Figure 5I right panel and S5). Hence, these data importantly suggest that DMF impacts the human intestinal microbiota in a way that appears individual specific, with potential consequences for microbiota-host interaction and intestinal homeostasis.

## DISCUSSION

Succination describes modification of Cys residues by fumarate and derivatives. Fumarate esters, DMF and MMF, are exploited as drugs against a wide diversity of diseases, from fungal infection to neurological disorders. Here, we report the results from a multi-leveled analysis of the impact of succination in prokaryotes. *E. coli* was used as a model for conducting biochemical, genetic and physiological analysis of exposure and deciphering molecular basis of DMF and MMF effects. Main cellular pathways targeted by DMF/MMF are related to sulfur metabolism, oxidative stress, proteostasis. with a strong effect on [Fe-S] cluster bound proteins. Furthermore, we showed that gut microbiota model OMM^12^ and human fecal communities cultivated in a MBRA system exhibited profound modification of their composition following exposure to MMF and DMF treatments.

Proteomic analysis of FAEs exposed *E. coli* revealed it contains a wide panel of Cys_suc_ containing proteins. Exposure to FAEs was associated with high abundance levels of chaperones and proteases, suggesting the cell system is trying to repair damaged proteins and solve folding related issues. Moreover, imaging-based techniques allowed us to visualize FAEs-caused formation of protein aggregates *in vivo* as observed using the IbpA-msfGFP biosensor. We concluded that FAEs are potential threats for protein homeostasis. This echoes studies in adipocytes that suggested that succination by fumarate inactivates the disulfide isomerase —a key protein for disulfide bond formation in endoplasmic reticulum— triggering misfolding stress ^26^.

The [Fe-S] proteins constituted a large fraction of those proteins showing lower abundance in the *E. coli* DMF-treated cells. Liganding of a vast majority of [Fe-S] clusters is mediated via Cys residues. Hence an immediate hypothesis was that Cys residues were succinated and lose their liganding capacity. However, such a view is at odds with our *in vitro* experiments with FumA, in which no loss in activity was observed when the [Fe-S] FumA form was exposed to FAE. This indicated that [Fe-S] clusters protected Cys residues from FAEs attack. In contrast, when apo-FumA was first treated with FAEs, [Fe-S] cluster insertion was impossible afterwards. The simplest explanation is that Cys residues being exposed in the apo-form of FumA, they have been succinated by FAEs and were eventually unable to bind an [Fe-S] cluster. Thus, to understand why [Fe-S] proteins were among the most altered by FAEs treatment, one needs to consider another effect that FAEs treatment imposes on *E. coli*, that is, oxidative stress. In particular AhpC was found to be succinated at its Cys_166_ residue which is essential for peroxide (H_2_O_2_) reduction ^27^, likely enhancing the level of accumulated intracellular H_2_O_2_. Moreover, expression of genes under the control of the H_2_O_2_-sensing OxyR regulator, including *ahpC*, were found to be induced by FAEs. Last, DMF exposure caused depletion of ROS-buffering GSH and GSSG another lead to enhanced intracellular ROS level. Hench, we propose that ROS will first destabilize exposed clusters of [Fe-S] proteins, increasing apoforms, whose exposed Cys residues are succinated by DMF. These succinated polypeptides cannot rebind [Fe-S] clusters and are eventually degraded, explaining their reduced abundance in DMF treated cells.

Another protein target of FAEs as revealed by proteomic analysis was GAPDH whose Cys_150_ catalytic residue was succinated. The GapA Cys_150_ was found in the three different succination species in all replicates (2SC, MSC and DSC). This is consistent with the strong nucleophilicity of GapA Cys_150_ ^28^. Interestingly, the solvent exposed AhpC Cys_166_ residue was also found in the three different succinated states as well. That such widely different proteins, GapA and AhpC, with different physico-chemical surroundings of the targeted Cys residues, reacted similarly showed how difficult it might be to predict potential Cys targets of DMF. As a matter of fact, previous studies have searched for signatures or patterns that could help predicting potential FAEs or fumarate targeted Cys residues but were quite unsuccessful ^16,17^. Here, a motif, reading ECxP was identified but found in only 3 peptides (in ArgE, Dcm and OxyR proteins), thus it cannot be used to identify/predict Cys_suc_. Overall, succinated sites often had at least one negative charged residue (D or E) within ± 2 residues, though this not always predicts high succination rates (e.g. GapA Cys_150_). Notably, this bacterial motif differs from eukaryotic models, which show enrichment of basic flanking residues ^17^.

DMF and MMF are active ingredients of routinely used drugs in clinics. These drugs are taken by oral route and it was of interest to test their effects, if any, on gut microbiota. Our study revealed that both DMF and MMF exert toxic effects on a wide range of mouse gut microbiota species, impacting both individual populations and mixed consortia. Notably, we found that MMF was more toxic than DMF. Additionally, we identified three microbiota species —*B. animalis* (an actinobacterium), *E. faecalis* and *L. reuteri* (both Firmicutes)— which exhibited immediate resistance when cultured in isolation with mM concentrations of FAEs. Interestingly, other firmicutes phyla members showed either immediate or delayed resistance after several hours of exposure to FAEs in single cultures. Several reasons can be put forward, such as low permeability of FAEs due to the membrane composition, activation of efflux pumps, resistance determinants to succination or detoxification pathways. Regarding the last possibility, it is worth mentioning that recent studies have identified two distinct evolutionary conserved pathways in firmicutes and α-proteobacteria that catabolize 2SC ^29,30^. The *yxeKLMNOPQ* operon characterized in *Bacillus subitilis* ^29^ was shown to use YxeK to breakdown 2SC adducts via oxygenolytic C–S bond cleavage ^31^. The second pathway employs a second type of S-(2-succino) lyase (2SL) that replaces *yxeK* ^30^. Neither YxeK nor 2SL homologs were found in the strains studied in the present work, indicating that additional new resistance determinants to FAEs remain to be identified.

The OMM^12^ model system has been widely used for studies regarding, among others, colonization resistance to pathogens, intestinal metabolic patterns and community interactions and structure ^24,32^. Our results showed that microbial community could provide protection to otherwise sensitive species, such as *E. coli*, *A. muciniphila*, *B. caecimuris*, *M. intestinale* and *T. muris*, which were vulnerable in isolated cultures. A. muciniphila is considered a promising next-generation beneficial microbe as it shows a higher prevalence in patients suffering from relapsing-remitting multiple sclerosis. Our results showed that *A. muciniphila* benefited from the OMM^12^ community to resist DMF toxicity.

Importantly, our MBRA experiments with a more complex microbiota community confirmed that FAEs could alter the gut microbiota composition with different or equal impact on different microbiota members. Interestingly, members of some Bacilli class seem to well adapt to FAEs exposure in a community set up or in isolated cultures, as evidenced for *Enterococcus* sp. and *Streptococcus* sp. in the MBRA and *E. faecalis* and *L. reuteri* of the OMM^12^ strains.

In conclusion our study provides a mechanistic understanding of FAE action in whole bacterial cells. It shows how gut communities contained species exhibited widely variable susceptibility to FAE, some species being highly tolerant. It will be important to test whether such a different fitness or protective effect holds in a whole animal, which has a different consortium composition and would integrate host response.

### Limitations of the study

While our study clearly demonstrates the effects of FAE on gut bacterial communities, we acknowledge that the rapid conversion of DMF to its active metabolite MMF in the human gut, along with pharmacokinetic factors such as pH-dependent hydrolysis and esterase activity, may result in bacterial exposure levels that differ from the conditions used here.

## Supporting information

SupTable_S1

SupTable_S2

SupTable_S3

SupTable_S4

SupTable_S5

## RESOURCE AVAILABILITY

### Lead contact

Further information and requests for resources and reagents should be directed to and will be fulfilled by the lead contact Rodrigo Arias-Cartin or to Frédéric Barras

### Materials availability

No unique materials were generated in this study.

### Data and code availability

The accession number for the mass spectrometry proteomics data reported in this paper is PRIDE: PXD062967. Code for Image analysis has been deposited at https://github.com/marieanselmet/DMF_MMF_microscopy_analysis/tree/main. Microscopy data reported in this paper was deposited at DOI:10.17632/rg7db6xyh7.1. Any additional information required to reanalyze the data reported in this paper is available from the lead contact upon request.

## ACKNOWLEDGMENTS

We wish to thank the SAMe Unit members and Melodi Birepinte for insightful comments. R.A.C thanks María José Navarro Porras for experimental support. M.A., F.B., J-M B. and R.A-C acknowledge support from Institut Pasteur, CNRS and Agence Nationale de la Recherche ANR-10-LABX-62-IBEID. A.G.B. thanks Diana Ring for experimental support. This work was partially supported by Microbiology Collaborative seed call Groot-21 FUMA project granted to R.A-C. We thank the Pasteur Institute Image Analysis Hub for advice with the microscopy image analysis.

## AUTHOR CONTRIBUTIONS

Writing original manuscript, F.B. and R.A-C.; writing, reviewing and editing, M.A., T.D., A.G.B., J-M.B., M.M., B.S. and B.C.; conceptualization and design, B.S., B.C., F.B and R.A-C; bioinformatic and microscopy analysis, M.A. and R.A-C; mass spectrometry experiments and analysis, T.D. and M.M.; microbial community experiments, A.G.B and E.B.; toxicity experiments on *E. coli*, B.J-A.; enzymatic experiments, J-M. B. and R.A-C.; *E. coli* strains construction R.A-C.; funding acquisition and supervision F.B. and R.A-C.

## DECLARATION OF GENERATIVE AI AND AI-ASSISTED TECHNOLOGIES

During the preparation of this work the authors used ChatGPT and Gemini for language polishing or drafting text. After using this tool, the authors reviewed and edited the content as needed and take full responsibility for the content of the published article.

## SUPPLEMENTAL INFORMATION TITLES AND LEGENDS

### SupPDF_Anselmetetal2025. Figures S1–S5

**Figure S1.**
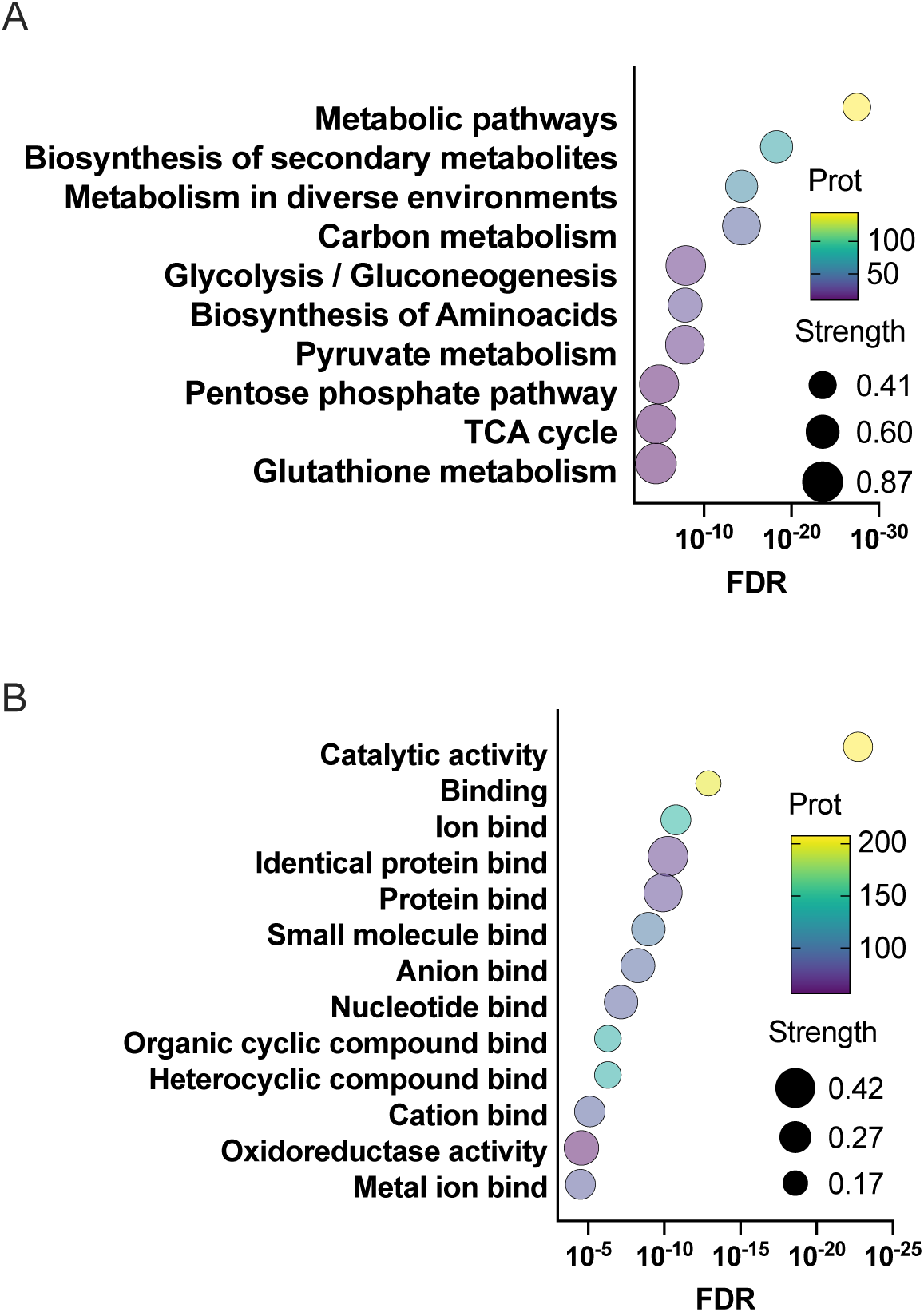
Enrichment analysis of the succinated proteins. Proteins were analyzed according to KEGG biological process (A) or GOterm molecular function (B). FDR: False Discovery Rate. Prot scale bar: color indicates the number of proteins within a particular category. Strength: Log_10_(observed /expected).

**Figure S2.**
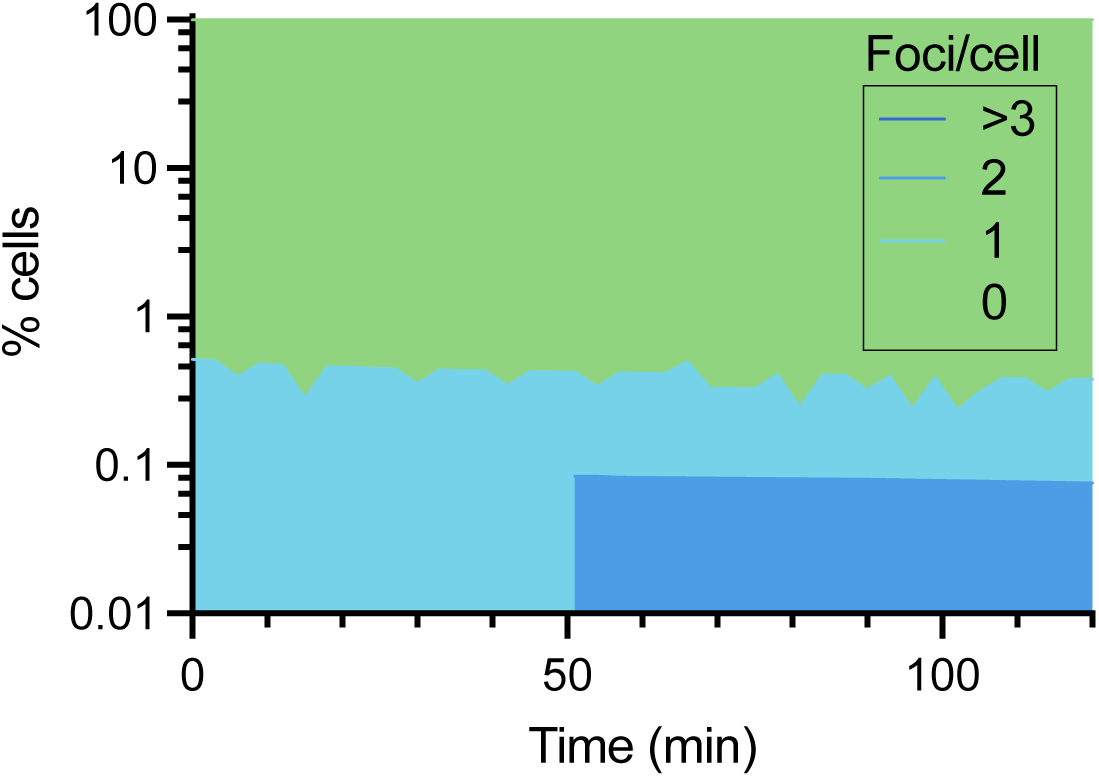
Analysis of time lapse experiment of cells producing IbpA-msfGFP foci upon DMSO exposure. Related to. Figure 3

**Figure S3.**
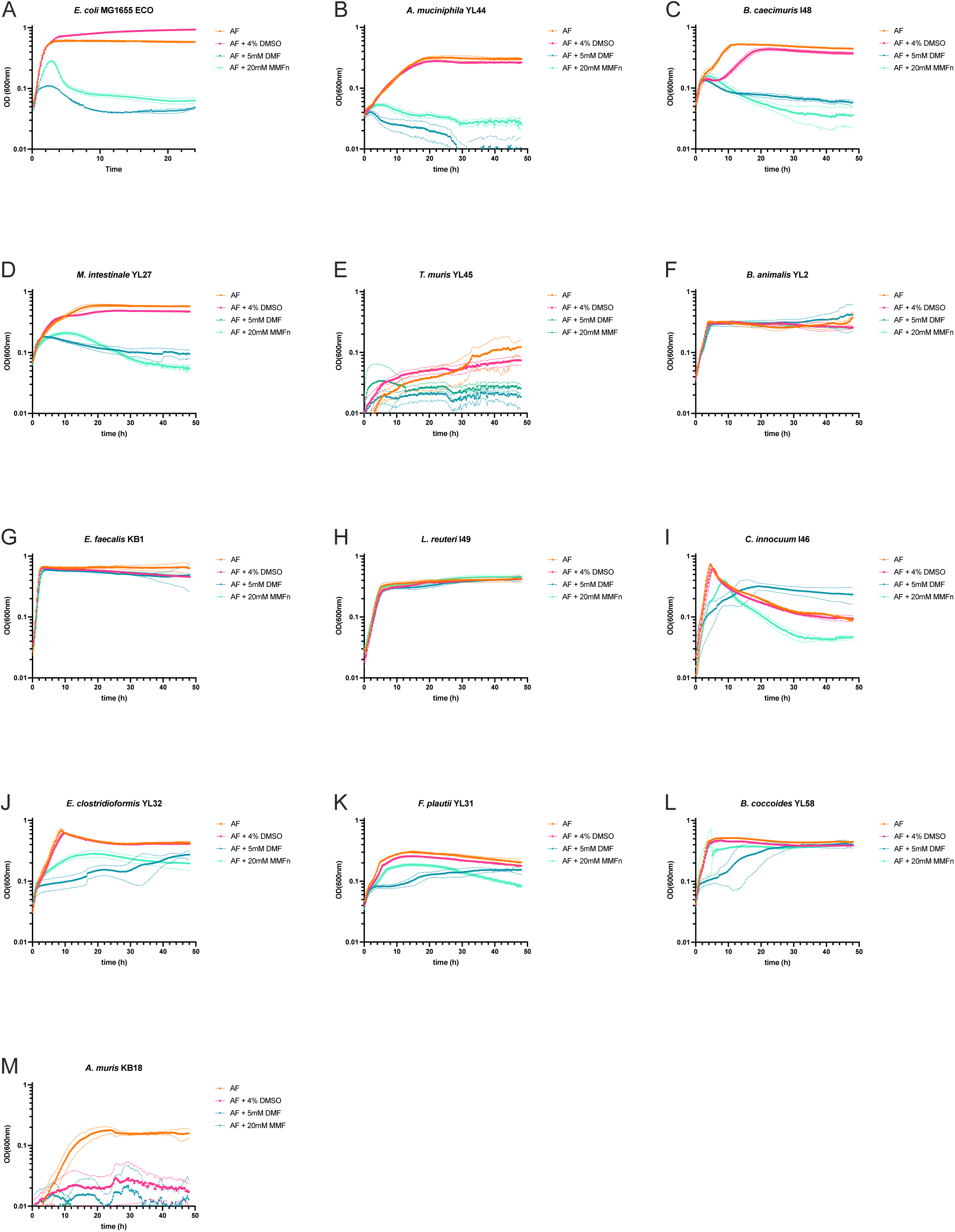
Effect of FAEs on OMM^12^ and *E. coli* strains isolated cultures, related to Figure 5. Representative anaerobic growth curves of each strain exposed to 5 mM DMF or 20 mM MMF in AF medium. (A) *E. coli* MG1655, (B) *A. municiphila*, (C) *B. caecimuris*, (D) *M. intestinale*, (E) *T. muris*, (F) *B. animalis*, (G) *E. faecalis*, (H) *L. reuteri*, (I) *C. innocuum*, (J) *E. clostridioformis*, (K) *F. plautii*, (L) *B. coccoides* and (M) *A. muris*. Medium pH readjusted to pH 7.0 after addition of MMF, except for (E) and (M). DMSO control was equivalent to the highest drug concentration tested (4% v/v). n = 3 per treatment. Standard deviation is represented by dotted lines.

**Figure S4.**
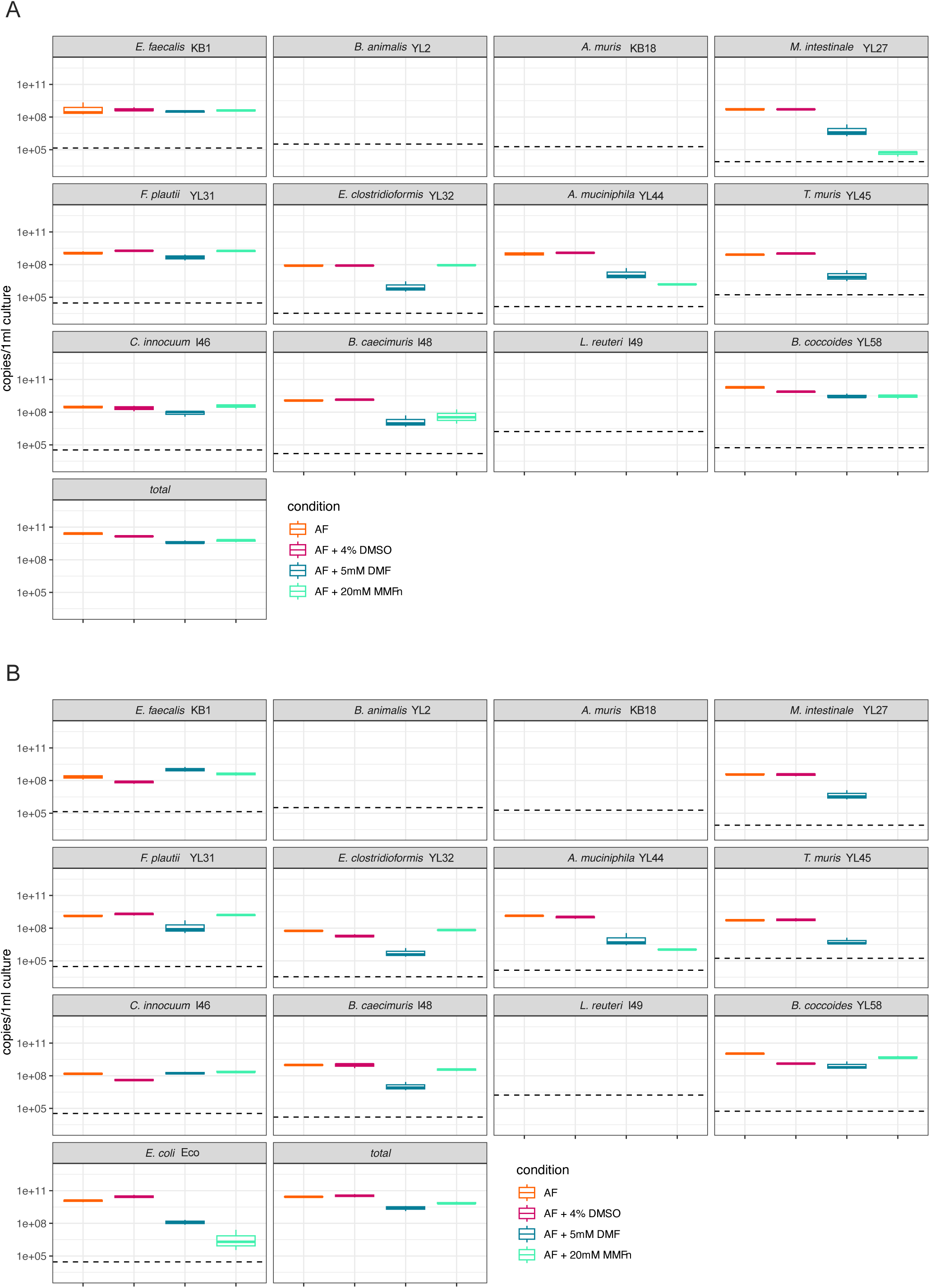
Effect of FAEs on OMM12 and OMM12+Eco consortiums, related to Figure 5. qPCR absolute analysis on consortium after 3 days exposure to (A) 5 mM DMF and (B) 20 mM MMF. DMSO control was equivalent to the highest drug concentration tested (4% v/v). n = 3 per treatment. Standard deviation is represented by upper and lower bars from average. Dotted line represents threshold limit.

**Figure S5.**
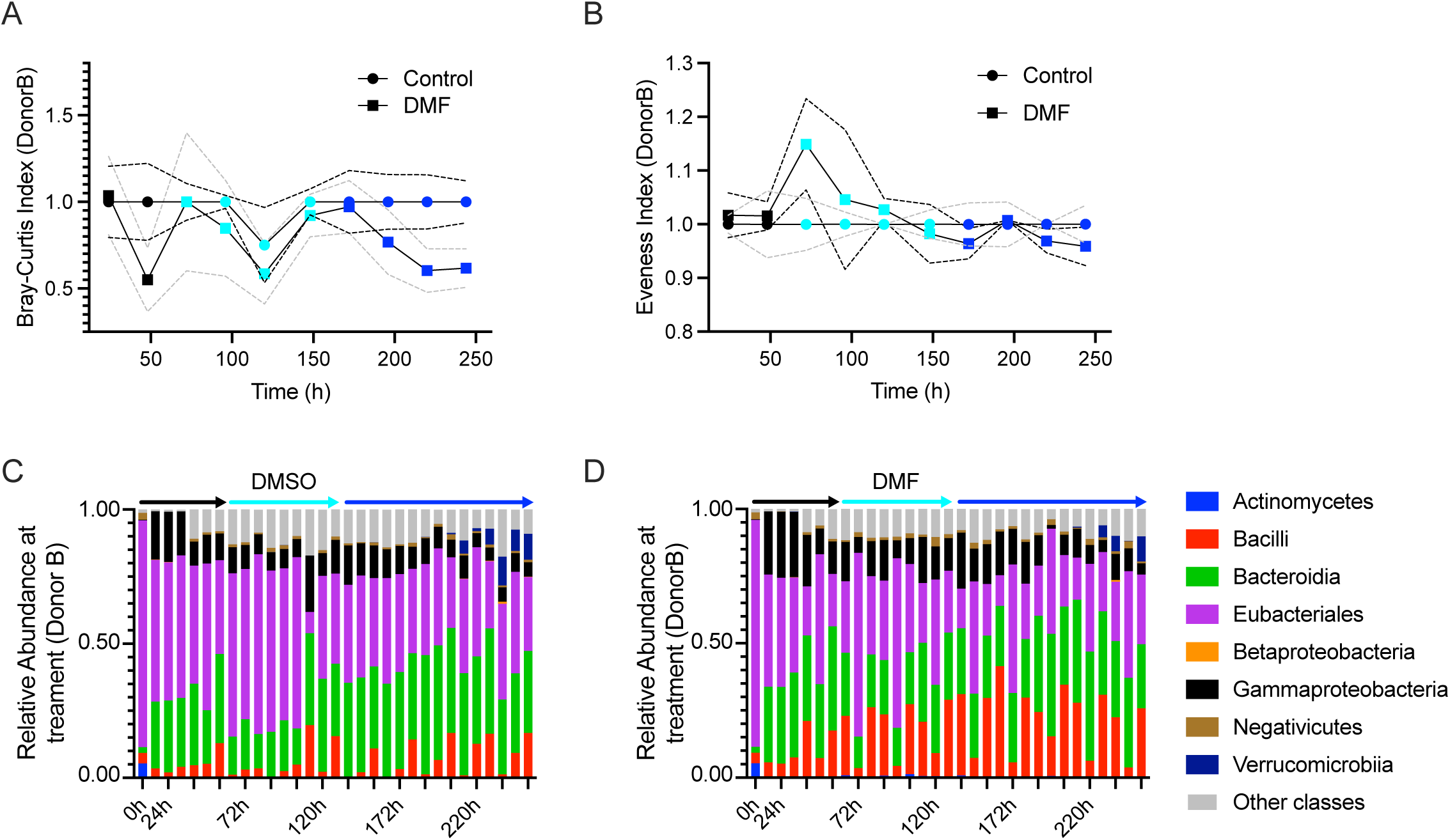
Dynamics of MBRA consortiums upon exposure to FAE in Donor B, related to. Figure 5. (A) Bray-Curtis distance and (B) Evenness index analysis of MBRA samples generated with the DMF and DMSO control from Donor A samples. Data were normalized to the DMSO control at each time point, defined as 1. Dots color indicates the stabilization, treatment and post-treatment points. Solid and dotted lines correspond to the mean and standard deviation of triplicates of each time point, respectively. (C) Relative abundance of the MBRA control and (D) DMF-exposed microbiota consortiums at Class taxonomic level during a 10-day long experiment. Abundance of each individual class is shown as relative abundance and expressed as percentage of cumulative 16S rRNA gene copy numbers of the rest of the classes. One bar corresponds to one of three replicates. Arrow color corresponds to the experiment section, stabilization (black), treatment (cyan) and posttreatment (blue).

**Table S1. Identification of succinated peptides in *E. coli* DMF-treated cultures using LC-MS/MS analysis. Excel file containing additional data too large to fit in a PDF, related to Figure 1**.

**Table S2. STRING-based functional enrichment analysis of succinated and differentially abundant proteins in DMF-treated *E. coli*. Excel file containing additional data too large to fit in a PDF, related to Figure 2 and Figure S1.**

**Table S3. Comparative analysis on DMF-treated *E. coli* vs DMSO-exposed *E. coli* using LC-MS/MS proteomics. Excel file containing additional data too large to fit in a PDF, related to Figure 2**.

**Table S4. Identification of succinated peptides on apo and holo FumA exposed to FAEs or DMSO control using LC-MS/MS analysis. Excel file containing additional data too large to fit in a PDF, related to Figure 4**.

**Table S5. qPCR raw and relative data from OMM**^12^ **and OMM^12+Eco^ experiments at Day 3. Excel file containing additional data too large to fit in a PDF, related to Figure 5**.

## STAR*METHODS

### KEY RESOURCES TABLE

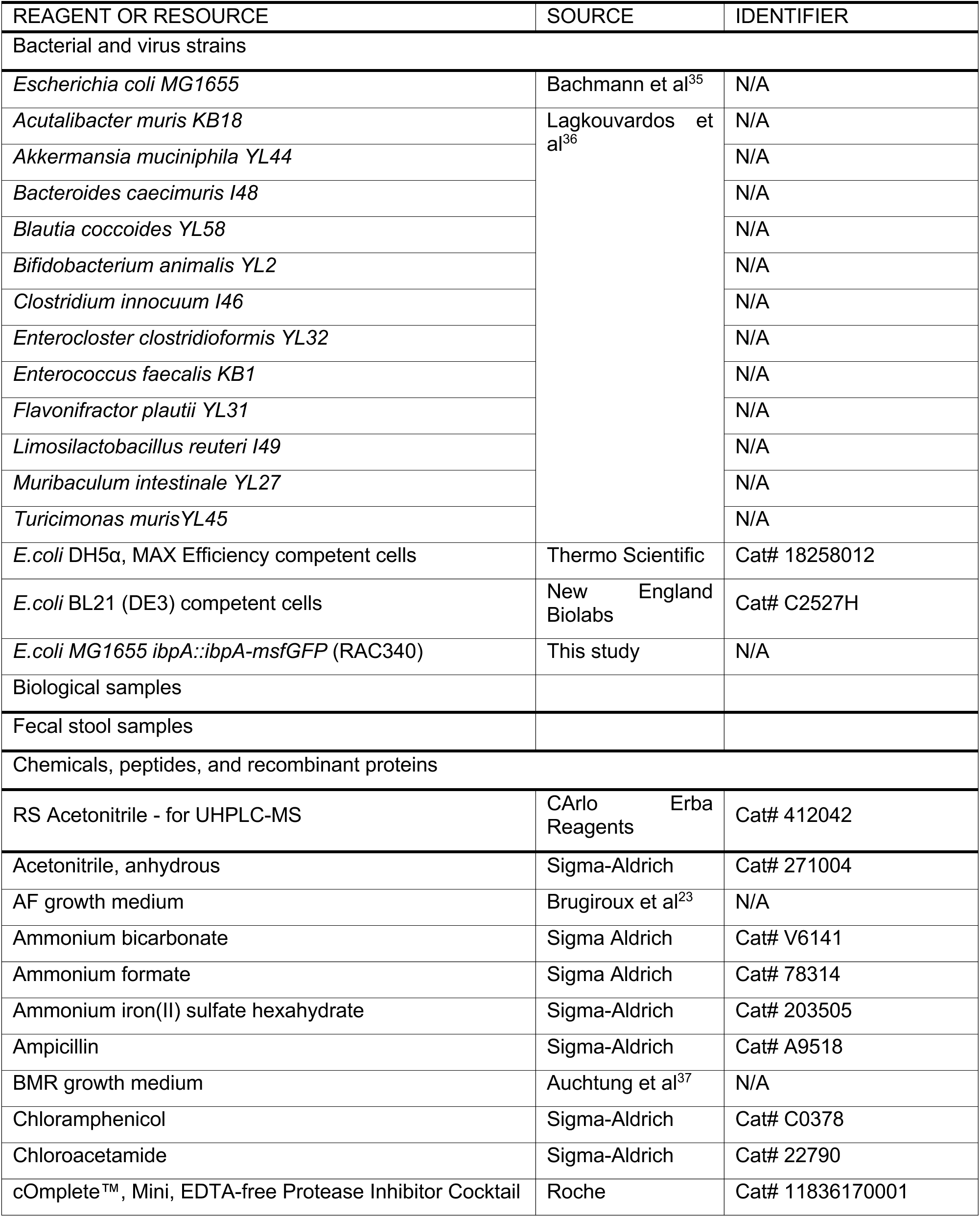
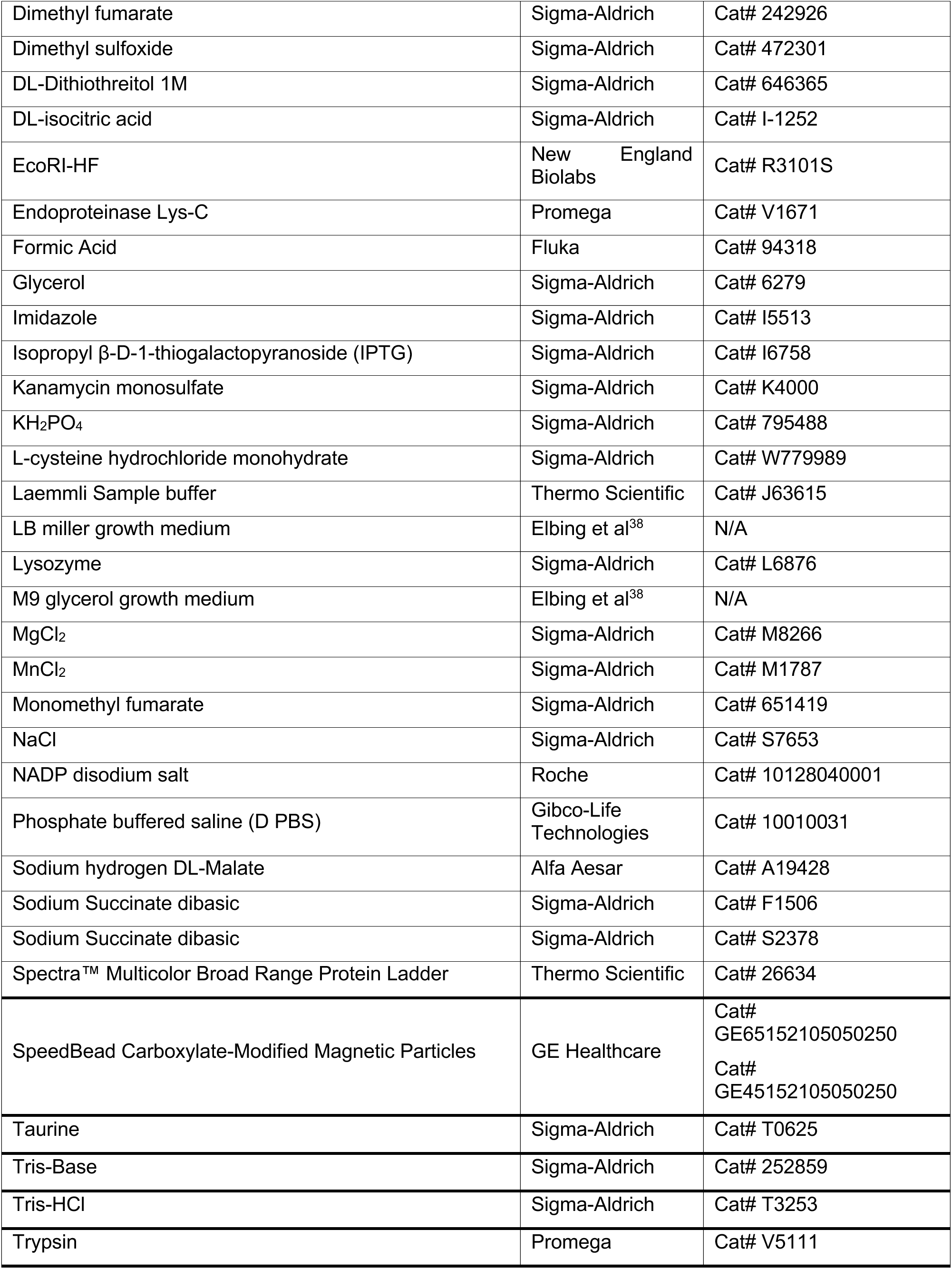
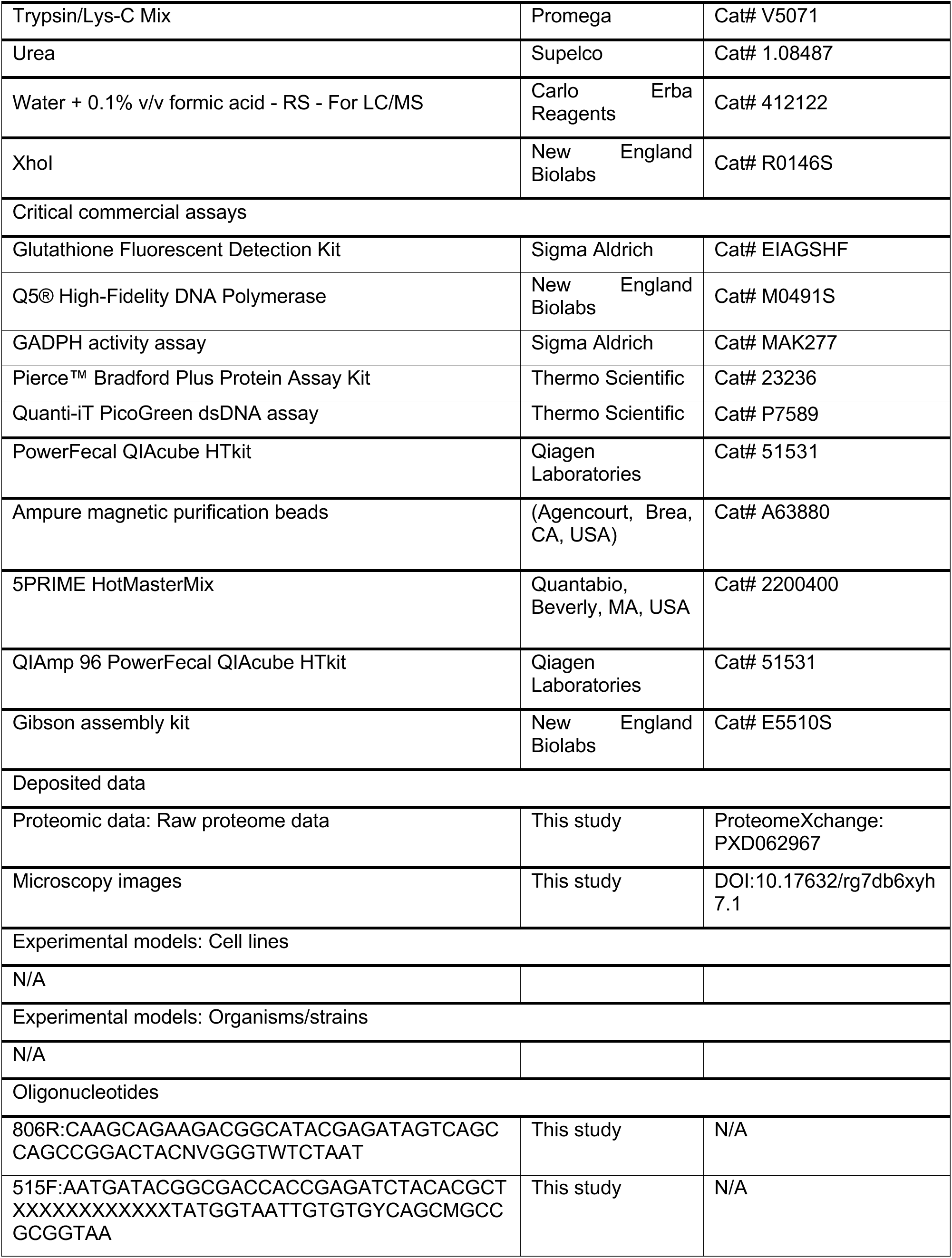
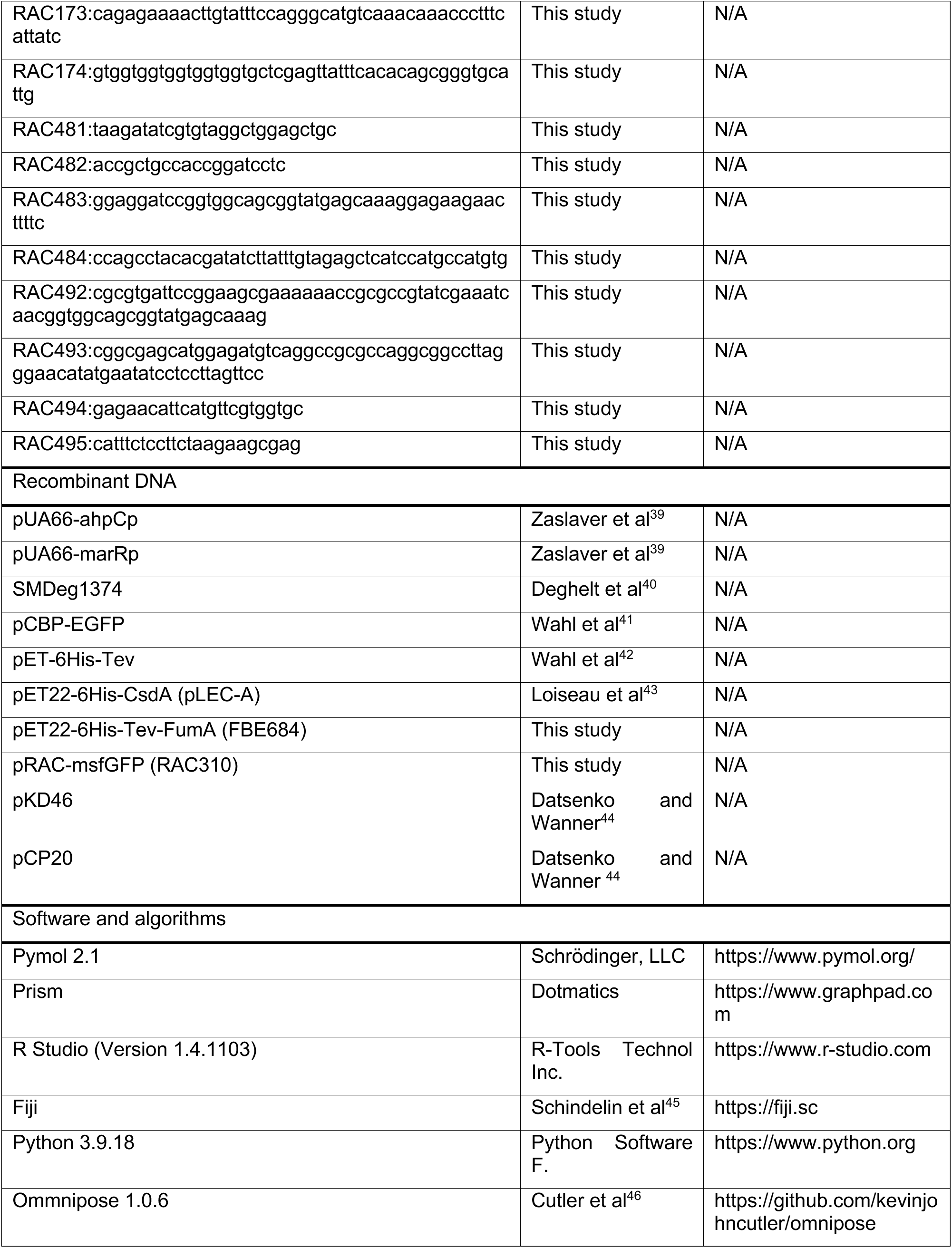
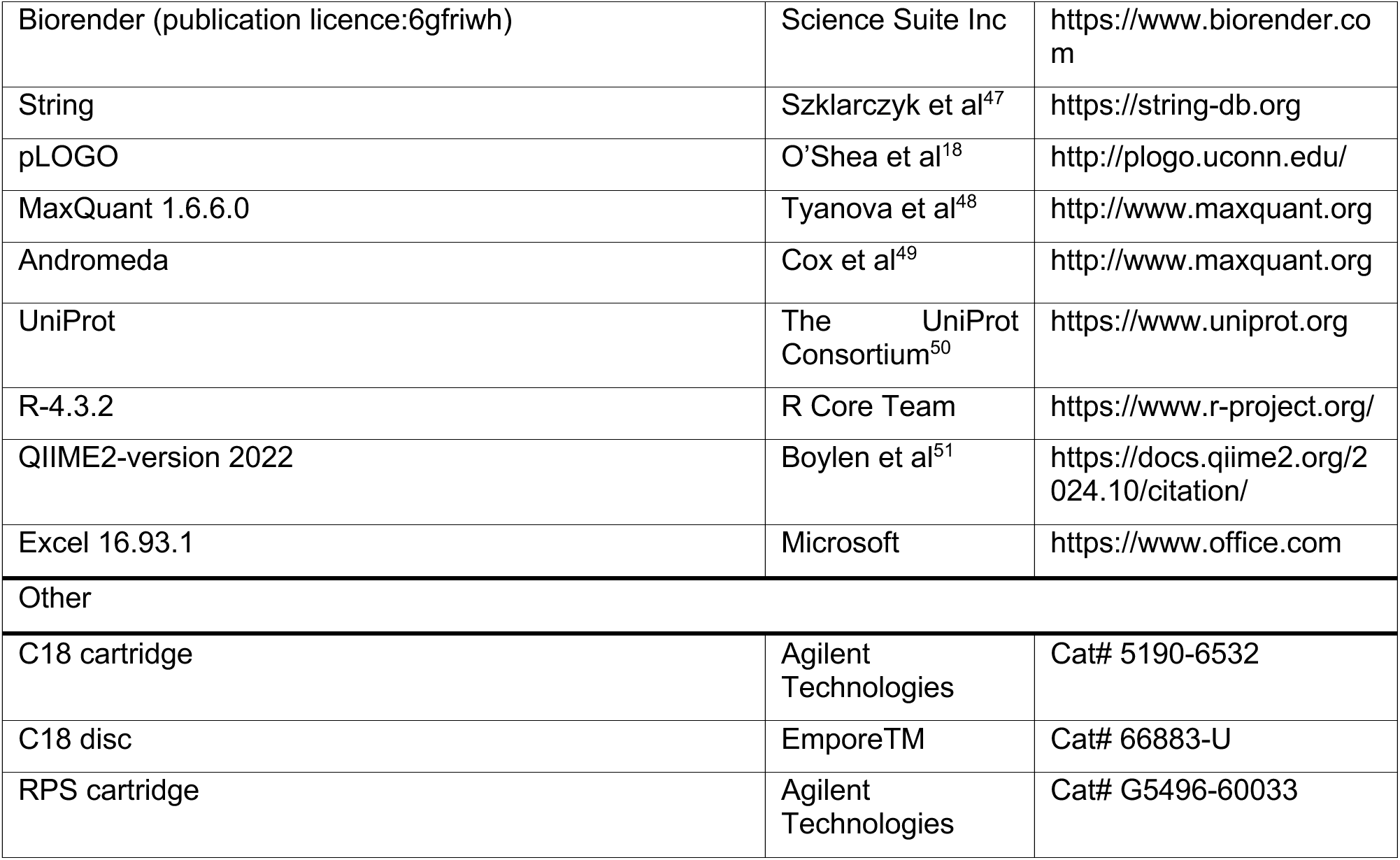

### EXPERIMENTAL MODEL AND STUDY PARTICIPANT DETAILS

Microbial experimental models used in this study are detailed in the reagents and source table.

## METHOD DETAILS

### Bacterial growth and reagents

All bacterial cultures were grown at 37°C and 200 rpm in LB Miller (5 gr·L NaCl adjusted to pH 7.3) or M9 medium. DMF and MMF stock solutions were prepared in DMSO at 500 mM and 200 mM, respectively and kept up to 3 months at room temperature. Working solutions were freshly prepared by diluting stock solutions in the appropriate medium at the desired concentration. When necessary, antibiotics were used at: Ampicillin 100 µg·mL, Chloramphenicol 25 µg·mL and Kanamycin 50 µg·mL.

### OMM^12^ and OMM^12+Eco^ experiments

OMM strains (indicated Reagents or resource table) were cultivated at 37°C without shaking in a Coy anoxic chamber (gas atmosphere 7% H_2_, 10% CO_2_, 83% N_2_) in AF medium (18 g·L^−1^ brain-heart infusion, 15 g·L^−1^ trypticase soy broth, 5 g·L^−1^ yeast extract, 2.5 g·L^−1^ K_2_HPO_4_, 1 mg·L^−1^ hemin, 0.5 mg·L^−1^ menadione, 3% heat-inactivated fetal calf serum, 0.25 g·L^−1^ cysteine- HCl‧H_2_O)^23^. All strains were prepared from frozen monoculture stocks and subcultured once before use for growth curves or inoculum preparation. Growth curves were recorded in 96 well format in a Biotek Epoch 2 (Agilent) microplate spectrophotometer inside an anoxic chamber. Community inocula were prepared by diluting all single strain subcultures to starting with an optical density at 600 nm (OD600) 0.1 with fresh medium and mixing in equal ratios. Communities were inoculated with 10% and further diluted 1:10 every 24 hours for 5 days for community equilibration. DMF, MMF and DMSO controls were added to the medium for the next three days. For MMF-neutralized condition, medium was readjusted to pH 7.0 after MMF addition using 5M KOH.

### DNA extraction and qPCR

DNA was extracted from cell pellets using phenol-chloroform extraction after bead beating following the protocol from Turnbaugh et al^52^ and further purified using a column-based gDNA purification kit (Macherey-Nagel). Resulting DNA was used to perform qPCR on a Roche Lightcycler 96 system with strain-specific primers and probes^23^. 5 ng of purified gDNA were used as template.

### MiniBioReactor Arrays (MBRAs) experiment

#### Fecal sample collection

Fecal samples were provided by two healthy volunteers and collected into sterile containers, sealed and transferred into an anaerobic chamber within 10 min of defecation. Here, fecal samples were manually homogenized and aliquoted into sterile 50 mL tubes and then stored at -80°C until use. The research protocol was approved by the GSU IRB committee under approval number H19174. Individuals donating samples provided informed consent prior to donation.

#### MBRA setup, experimental plan and timepoints for samples collection

MBRA systems were prepared as previously described^53^ and housed in an anaerobic chamber. The system consists in 24 chambers filled with 15 mL of Bioreactor Medium (BRM). Chambers were connected to two 24-channel peristaltic pumps with low flow-rate capabilities (205S peristaltic pump with 24-channel drive, Watson-Marlow) and held on a magnetic stand for continual homogenization of the medium. Following autoclaving, MBRA chambers, tubing, and the BRM medium were placed in the anaerobic chambers for at least 72 h. MBRA chambers were subsequently filled with BRM and inoculated with the fecal sample. For the inoculation, fecal samples were resuspended at 10% w/v in anaerobic phosphate buffered saline (D PBS) in the anaerobic chamber, vortexed for 5 min and centrifuged at 800 rpm for 5 min at 20°C. Supernatants were collected in the anaerobic chamber and filtered through a 100 µm filter in order to remove any particles. The inoculation volume of the fecal slurry was set 3.8 mL per chamber ^53^. After inoculation, fecal bacterial communities were allowed to equilibrate for 16 h prior to flow initiation at 1.875 mL·h (8h retention time). As presented in Figure 5D, D-1 timepoint corresponds to the inoculation with fecal slurry, and D3 (72h) timepoint corresponds to the first injection of DMF (reference for DMF). A second and a third injection were performed at D4 (96h) and D5 (120h) and post-treatment periods lasted until D10. Starting from D0, 400 µL of samples were collected at various time points (Figure 5D) and collected samples were stored at -80°C until analysis.

#### Bacterial DNA extraction

DNA was extracted from frozen MBRA suspension or fecal samples using a QIAmp 96 PowerFecal QIAcube HTkit with mechanical disruption. Briefly, 650 µL of prewarmed buffer PW1 were added to the samples. Subsequently, samples were thoroughly homogenized using bead-beating with a TissueLyser before centrifuging the plate at 4000 rpm for 5 min at 20°C to pellet beads the particles. Four hundred microliters of supernatant were transferred to a new 96-well plate containing 150 µL of buffer C3. After mixing and incubation on ice for 5 min, centrifugation was performed at 4000 rpm for 5 min at 20°C. Three hundred microliters of each supernatant were then transferred to a new 96-well S-block plate and 20 µL of Proteinase K were added and incubated for 10 min at room temperature. The following steps were performed on a QIAcube high-throughput robot as follow: addition of 500 µL of Buffer C4, DNA binding to a QIAmp 96 plate, column wash using 800 µL of AW1, 600 µL of AW2 and 400 µL of ethanol, and elution by adding 100 µL of ATE buffer.

#### Microbiota analysis through 16S rRNA gene sequencing

16S rRNA gene amplification and sequencing were performed using the Illumina MiSeq technology following the protocol of the Earth Microbiome Project^54^, with some modifications. Briefly, the 16S rRNA genes, region V4, were PCR amplified from each sample using a composite forward and reverse primer containing a unique 12-base barcode, designed with the Golay error-correcting scheme used to tag PCR products from respective samples^54^. PCR reactions consisted of 5PRIME HotMasterMix 0.2 µM of each primer (515F and 806R), 10-100 ng template, and reaction conditions were set as follow: 3 min at 95°C, followed by 30 cycles of 45 s at 95°C, 60 s at 50°C and 90 s at 72°C on a Biorad thermocycler. PCR products were then visualized by gel electrophoresis and quantified using Quanti-iT PicoGreen dsDNA assay. A master DNA pool was generated from the purified products in equimolar ratios and subsequently purified with Ampure magnetic purification beads. The obtained purified pool was quantified with the Quanti-iT PicoGreen dsDNA assay, followed by sequencing using an Illumina MiSeq sequencer (pair-end reads, 2 x 250 bp) at the GENOM’IC core facility at Cochin Institut, Paris, France.

#### 16S rRNA gene sequence analysis

QIIME2-version 2022 was used to analyze 16S rRNA sequences^51^. These sequences were demultiplexed and quality filtered using Dada2 method 3 with QIIME2 default parameters to detect and correct Illumina amplicon sequence data, generating a table of Qiime 2 artifact. Then, a tree was generated using the align-to-tree-mafft-fasttree command for phylogenetic diversity analysis and we computed alpha and beta diversity analyses using the core-metrics-phylogenetic command. Principal Coordinate Analysis (PCoA) plots were used to assess variations between experimental groups (beta diversity). Alpha diversity was computed with the Evenness index. For the taxonomic analyses, features were assigned to operational taxonomic units (OTUs) with a 99% threshold of pairwise identity to the Greengenes reference database 13_8 4.

### Bacterial Strains and Plasmids

The *E. coli* plasmids and strains are described in Reagents or source table. Constructions were made as following: pET22-6Histev-FumA: *fumA* was amplified from *E. coli* MG1655 gDNA using primers RAC173 and RAC174. PCR fragment was introduced using Gibson assembly kit into a pre-digested pEB1188 at EcoRI-XhoI sites. pRACpmsfGFP: *msfGFP* was first amplified from SMDeg1374 using primers RAC483 and RAC484 an introduced using Gibson assembly kit into pCBP-EGFP amplified with primers RAC481 and RAC482. MG1655 *ibpA*::ibpA-msfGFP: *msfGFP* was introduced at *ibpA* 3’end using the Red recombinase Datsenko-Wanner system^44^. Briefly, *msfGFP* was amplified with primers to generate fragment with kanamycin (Kn) resistance marker flanked by FRT sites using primers RAC492 and RAC493 and pRACpmsfGFP as template. PCR fragment was electroporated into MG1655 harboring pKD46 and selected on LB Kn. The Kn cassette was later excised using pCP20 plasmid. Chromosomal fusion at *ibpA* was confirmed with primers 494-495.

### Determination of MMF and DMF Minimal inhibitory concentration and combinatorial toxicity

The inhibitory concentrations of DMF and MMF were determined as the lowest concentration required for growth inhibition of a microorganism after 12 to 16h of incubation in the desired medium at 37 °C with shaking in a 96-well plate and starting with OD600 0.01 from an overnight liquid culture. DMF was tested at concentrations of 5 mM, 2.5 mM, 1.25 mM, 0.625 mM and 0.3125 mM, while MMF was tested 20 mM, 15 mM, 10 mM, 5 mM, 2.5 mM and 1.25 mM. Each concentration was assayed in triplicates in 200 µL medium and incubated at 37 °C and 800 rpm linear shaking using a TECAN Spark Microplate Reader at 37°C. Follow-up combinatorial chemical complementation was done by adding 0.5 mM, 0.05 mM and 0.005 mM of L-cysteine or 5.0 mM 0.5 mM, 0.05 mM and 0.005 mM of Taurine from 20 mM and 25 mM stock solutions, respectively.

### Enzymatic assays

*E. coli* MG1655 lysates cells were cultured in LB medium and exposed for 1 h 30 min to MMF, DMF or DMSO starting at exponential growth phase (OD600 0.4-0.6) at 37°C and agitation. Then cells were harvested by centrifugation and lysed after spheroplast preparation using a modified Osborn et al^55^ protocol. Cell pellets normalized to the same OD600 were suspended in 10 mM Tris-HCl (pH 7.5) containing 0.7 M sucrose. Lysozyme (0.2 mg·mL^−1^) and EDTA (10 mM) were added, and the suspensions were incubated for 20 min at 4°C. The samples were centrifuged for 5 min at 10,000 rpm in an Eppendorf microcentrifuge and the supernatants were removed. The pellets were freeze-thawed, resuspended in cold water, and then sonicated for 10 sec with a Branson digital sonifier. Immediately after, a cuvette was loaded with 50 μL of cell lysate and 3 mL of buffer according to the enzyme to be tested and specific absorbance was monitored using Jasco V-730 spectrophotometer at 30°C. The aconitase activity was measured spectrophotometrically by monitoring the formation of cis-aconitate from isocitrate at 240 nm (ε = 3.6 mM−1·cm−1) in 50 mM Tris-HCl (pH 7.4) containing 20 mM isocitrate and 0.5 mM MnCl_2_ to initiate the reaction for cis-aconitate^56^. The fumarase activity was performed by monitoring the conversion of 50 mM L-malate to Fumarate at 250 nm (ε = 1.62 mM−1·cm−1) in 100 mM KH_2_PO_4_ at pH 7.6 using^57^. The succinate dehydrogenase activity was obtained by a coupled enzymatic test described by Jones et al 2013^58^ following the oxidation of NADPH concentration using the activity of at 340nm (ε = 4.81 mM−1·cm−1) on an AvaSpec-spectrophotometer. GADPH activity was obtained using the GAPDH Activity assay kit from Sigma by absorbance at 450 nm in a TECAN Spark Microplate Reader and following provider manual instructions. All specific activities were based on total protein content measured by the Bradford assay.

### Glutathione quantification

GSH and GSSG concentration was obtained using the Glutathione Fluorescent Detection Kit from Sigma (EIAGSHF) following provider manual instructions and standardized by mg of cells using a TECAN Spark Microplate Reader.

### Fumarase A and Cysteine desulfurase A expression and purification

Plasmid pET22-6Histev-FumA was transformed into *E. coli* BL21(DE3) and cultures were grown in LB Miller medium containing 100 µg·ml ampicillin at 37°C and 200 rpm. When culture reached OD600 0.6, FumA expression was induced for 4h by the addition of 500 µM isopropyl-β-D-thiogalactoside (IPTG). Next, cells were harvested by centrifugation at 4°C 4000 xg for 10 min and the pellet was washed once with 50 mL PBS and kept at -80°C. From this point all the following steps were performed on ice or at 4°C. The pellet was resuspended in 40 mL Buffer A (Tris 25 mM NaCl 100 mM Imidazole 25 mM pH 7.6) and supplemented with 1 pill of cOmplete™ Mini EDTA-free Protease Inhibitor Cocktail, 5 mM MgCl_2_ and 300 µl of Benzonase. Cells were broken by one passage on a Cell Disruptor (Constant systems©) at 20000 psi and cell lysate was cleared by 20 min centrifugation at 11000 rpm into a 5804R centrifuge (Eppendorf). Supernatant was applied to a 5 mL HisTrap™ High Performance column (Cytiva) using an ÄKTA prime plus FPLC system and washed with 5 column volumes with BufferA. Protein was eluted using a 20 mL 0-100% gradient with Buffer B (Tris 25 mM NaCl 100 mM Imidazole 300 mM, glycerol 10% pH 7.6) at 1 mL·min. Fractions containing a peak at 280nm absorbance were collected and assessed by SDS-PAGE gel electrophoresis. Purest samples were pooled, DTT added at 1 mM final concentration was added to the total volume, transferred into a dialysis bag of 12-14000 MWCO and dialyzed against 2 L of Buffer D (Tris 50 mM NaCl 100 mM 100 ug·mL His-tagged TEV protease pH 7.8) at 16°C with constant stirring. TEV digested FumA was obtained by passage into a column containing 2 mL of Ni-NTA beads (Quiagen). Flow through was collected by gravity was concentrated and dialyzed in a 15 mL Amicon 30KDa (Millipore) by 10 min centrifugation at 4000 rpm in a 5804R. During spinning, volume in the Amicon was resuspend every 2min with a pipette to avoid concentration at the bottom and precipitation. Buffer exchange was achieved by three washes with 10-12 mL each with Buffer E (Tris-HCl 40 mM pH 7,8 NaCl 80 mM glycerol 20%) during 30 min spin. Protein content was determined by the Bradford assay and aliquots were kept at -80°C until use. For *E. coli* CsdA, protein expression and purification protocol using pET22-6His-CsdA plasmid was identical to the FumA protocol with the following modifications: Buffer B was Tris-HCl 25 mM NaCl 100 mM Imidazole 300 mM, glycerol 10% pH 7.6, Buffer D was Tris 50 mM NaCl 500 mM pH 7.8 and Buffer E was Tris-HCl 40 mM, DTT 1 mM NaCl 80 mM glycerol 10% pH 7.8. The His-Tag removal step was omitted for CsdA.

### Fumarase A activity *in vitro* experiments

Effect of FAE on purified FumA was conducted anaerobically at room temperature (23-25°C) in an JACOMEX Campus glove box containing less than 0.3 ppm O_2_, equipped with an AvaSpec-ULS2048CL-EVO spectrophotometer and connected to an Avanlight-DH-S-BAL light source. The effect of DMSO, DMF or MMF on FumA activity was tested by incubating 1 nmol FAE per FumA cysteine residue (1:9 nmol ratio) for 2h in Buffer R (Tris 50 mM NaCl 100 mM DTT 5 mM glycerol 5% pH 8) with pre or post reconstituted [Fe-S] cluster FumA. The [Fe-S] cluster reconstitution was conducted by treatment of 100 μM of protein with a 5-fold molar excess of both ferrous ammonium sulfate and L-cysteine in 1 mL Buffer R. The reaction was initiated by the addition of 2:100 CsdA:FumA molar equivalents and monitored by UV-visible absorption spectroscopy until saturation of absorbance at 410-420 nm. The protein was subsequently purified in an His Spintrap™ column (Cytiva) previously equilibrated with Buffer R without DTT. Column was centrifuged for 1 min at 800 rpm, flow through was collected and protein concentration quantified using Bradford assay. For Mass spectrometry analysis 3 experiment replicates were pooled and analysed as described in Sample preparation for proteomics, HPRP fractionation and LC-MS/MS analysis following sections

### Cell lysate preparation for proteomics

Analysis was performed in 5 different replicates for each condition using lysates prepared from *E. coli* MG1655 wild-type strain grown in LB medium. A 500 mL culture was inoculated at OD600 0.01 from an overnight culture until it reached 0.5-0.6 units. At this point culture was divided in two 250 mL cultures to either to 2.5 mM DMF or its DMSO volume equivalent for 2h under agitation. Then 50 mL of cells were collected by 10min centrifugation at 4000 rpm in a pre-cooled centrifuge. Supernatant was discarded and cell pellets were freeze-thawed and resuspended in cold 10 mL 25 mM Tris-HCl (pH 7.8) supplemented with cOmplete™ Mini EDTA-free Protease Inhibitor Cocktail and disrupted by one passage at 2 bars in a Constant Cell CF1 (Disruption Systems) at 4°C. Protein content was determined by the Bradford assay.

### Sample preparation for proteomics

In-solution digestion for *E. coli* proteome analysis was carried out as follows. For each sample, 80 µg total protein were homogenized until 65 µL final volume with 25 mM Tris-HCl, pH 8.0. Then, one volume of 8M urea were added before a reduction of disulfide bonds in 20 mM DTT final and alkylation of free cysteine in 55 mM chloroacetamide (CAA) final. Digestion was performed using Endoproteinase Lys-C at ratio 1:80 (enzyme : protein) during 4h at RT and then using trypsin at ratio 1:50 for 12h at 37°C before stopping the digestion with 1% Formic Acid (FA) final. Digested peptides were desalted with the AssayMAP Bravo (Agilent Technologies) using 5 μL bead volume C18 cartridge and eluted with 80% acetonitrile (ACN), 0.1% FA. Finally, the peptide solutions were speed-vac dried before high-pH reversed-phase (HPRP) fractionation.

SP3 digestion for FumA succination analysis was carried out as follows. Purified FumA samples were homogenized until 40 µL final volume in 1% SDS, 25 mM Tris-HCl (pH 8). After, samples were reduced and alkylated as previously described. Protein isolation and digestion were carried out using the Single-Pot Solid-Phase-enhanced Sample Preparation (SP3) method, with minor modifications from the original protocol as described by Hughes et al.^59^ In brief, SP3 beads were prepared by mixing hydrophilic and hydrophobic Sera-Mag SpeedBeads in a 1:1 (v/v) ratio. Then, 1 µL of beads were added to samples followed by the addition of anhydrous ACN to a final concentration of 75% (v/v). After 30 min at room temperature under agitation and 1 min on a magnet, the supernatant was removed before 2 washes with 80% ACN and one with 100% ACN. For digestion, 50 mM Tris-HCl (pH 8.0) containing Trypsin/Lys-C Mix was added to the bead-bound proteins at a protein-to-enzyme ratio of 20:1. The samples were incubated for 12 h at 37°C. After digestion, the supernatant containing peptides was collected into a new tube following 1 min on a magnet. The beads were then washed with water, mixed for 10 min at room temperature, and the supernatant was pooled with the initial peptide solution. The resulting peptides were acidified with FA to a final concentration of 1%. Peptides were desalted with the AssayMAP Bravo (Agilent Technologies) as previously described.

### HPRP fractionation

We proceeded to a peptide fractionation using the Fractionation v1.1 protocol of the AssayMAP Bravo (Agilent Technologies). The dried pooled sample (80 μg) was resuspended in 20 mM ammonium formate (AmF) buffer, pH 10 before a HPRP fractionation. RPS cartridge (5 μL bead volume) were primed with 100 μL 80% ACN, 0.1% FA and equilibrated with 100 μL 20 mM AmF, pH 10. The samples were loaded at 5 μL·min followed by an internal cartridge wash and cup wash with 50µL of 20 mM AmF, pH 10 at 5 μL·min. Elution steps were performed with 100 μL of 20%, 30%, 40% and 80% ACN in 20 mM AmF, pH 10 at 5 μL·min. A preexisting volume of 20 μL containing the same elution buffer was present in the collection plates upon elution. All fractions were speed-vac dried and resuspended in 2% ACN, 0.1% FA before injection.

### LC-MS/MS analysis

A nanochromatographic system Proxeon EASY-nLC 1200 (Thermo Fisher Scientific) was coupled on-line to a Q Exactive^TM^ Plus Mass Spectrometer (Thermo Fisher Scientific) using an integrated column oven (PRSO-V1 - Sonation GmbH, Biberach, germany). For each sample, peptides were loaded into a capillary column picotip silica emitter tip (home-made column, 40cm x 75 µm ID, 1.9 μm particles, 100 Å pore size, Reprosil-Pur Basic C18-HD resin, Dr. Maisch GmbH, Ammerbuch-Entringen, Germany) after an equilibration step in 100 % buffer A (H_2_O, 0.1 % FA). SP3 digestion: Peptides were eluted with a multi-step gradient from 2 to 7 % buffer B (80 % ACN, 0.1 % FA) in 2.5 min, 7 to 23 % in 35 min, 23 to 45 % in 15 min and 45 to 95 % in 5 min at a flow rate of 250 nL·min over 64.5 min. In-solution digestion: Peptides were eluted with a multi-step gradient from 2 to 7 % buffer B (80 % ACN, 0.1 % FA) in 5 min, 7 to 23 % in 70 min, 23 to 45 % in 30 min and 45 to 95 % in 5 min at a flow rate of 250 nL·min over 117 min. Column temperature was set to 60°C. MS data were acquired using Xcalibur software using a data-dependent Top 10 method with a survey scans (300-1700 m/z) at a resolution of 70,000 and a MS/MS scans (fixed first mass 100 m/z) at a resolution of 17,500. The AGC target and maximum injection time for the survey scans and the MS/MS scans were set to 3E6, 20 ms and 1E6, 60 ms (100 ms for FumA samples) respectively. The isolation window was set to 1.6 m/z and normalized collision energy fixed to 28 for HCD fragmentation. We used a minimum AGC target of 1E4 for an intensity threshold of 1.7E5 (1.0E5 for Fum A samples). Unassigned precursor ion charge states as well as 1, 7, 8 and >8 charged states were rejected and peptide match was disable. Exclude isotopes was enabled and selected ions were dynamically excluded for 20 seconds (FumA samples) or 45 seconds (in-solution digestion).

### Data Processing for protein identification

Raw data were analyzed using MaxQuant using the Andromeda. The MS/MS spectra were searched against a UniProt *Escherichia coli* strain K12 database (download in 27/11/2028 - 4,347 entries) and FumA modified sequence. Usual known mass spectrometry contaminants and reversed sequences of all entries were included, cysteine carbamidomethylation (+57.0215Da) was considered as fixed modification and cysteine succinations as dynamic modifications: S-(2-succinyl)cysteine (2SC +116.0110 Da), S-(2-monomethylsuccinyl) cysteine (MSC +130.0266 Da) and S-(2-dimethylsuccinyl) cysteine (DSC +144.0423 Da). Methionine oxidation and acetyl protein N-terminus were also considered as dynamic modification. As recommended in Merkley et al.^60^, succinations were recalculated in monoisotopic mass such as (mass_monoiso_ succination - mass_monoiso_ carbamidomethyl). Resulting recalculation were: 2SC + 58.9895 Da, MSC +73.0051 Da and DSC +87.0208 Da. Trypsin with a maximum of 3 missed cleavages were set for the search. The minimum peptide length was set to 7 amino acids and one unique peptide to the protein group was required for the protein identification. Second peptide was enabled to identify co-fragmentation events. The “match between runs” feature was applied for samples having the same experimental condition with a maximal retention time window of 0.7 minute. The main search peptide tolerance was set to 4.5 ppm and to 20 ppm for the MS/MS match tolerance. The false discovery rate (FDR) for peptide and protein identification was set to 0.01. Unique and razor peptides, with at least 2 ratio count were accepted for quantification. Quantification was performed using the XIC-based LFQ algorithm with the Fast LFQ mode as described in Cox et al^61^. The mass spectrometry proteomics data have been deposited to the ProteomeXchange Consortium via the PRIDE^62^ partner repository with the dataset identifier PXD062967.

### Statistical analyses

To find the proteins more abundant in one condition than in another, the LFQ intensities quantified using MaxQuant were compared. Only proteins with at least 4 quantified intensity values in one of the two compared conditions were kept for further statistics. Proteins absent in a condition and present in another are put aside. These proteins can directly be assumed differentially abundant between the conditions. After this filtering, intensities of the remaining proteins were first log-transformed (log2) and normalized by centering the mean of the medians of intensities in each sample of the condition. Statistical testing was conducted using a limma t-test thanks to the R package limma^63^. An adaptive Benjamini-Hochberg procedure was applied on the resulting p-values thanks to the function adjust.p of the cp4p R package^64^. We considered as differentially abundant proteins those exclusive or absent in one or the other condition and those with an adjusted p-value inferior to a FDR (false discovery rate) level of 1% and an absolute log_2_ (fold-change) superior to |1|. Finally, the proteins of interest are therefore those which emerge from this statistical analysis supplemented by those which are considered to be present from one condition and absent in another.

### Microscopy experiments

Images were acquired with a fully motorized Nikon Ti-2E Eclipse inverted microscope equipped with a CFI Plan Apochromat λ DM 100XH 1.45/0.13 mm Ph3 oil objective (Nikon), a SpectraX illuminator (Lumencor), an Orca-Flash 4.0 V3 camera (Hamamatsu) and a GFP-1828A cube filtre (Ex: 482/18nm, Dichroic mirror: 495nm and BA: 500/24nm). Multi-dimensional image acquisition was made using NIS-Ar software (Nikon).The IbpA-msfGFP producing strain was cultured at 37°C in M9 0.2% glucose CAA 0.1% medium until OD600 0.5 using a 1/1000 dilution from an overnight preculture. Cells were immobilized in same medium supplemented with 1% agarose and imaged using 80ms phase contrast (ph) and 200 ms GFP acquisition time. For MG1655 pUA66-*ahpCp* and MG1655 pUA66-*marRp,* cells were cultured at 37°C in M9 0.4% glycerol medium supplemented with kanamycin until OD600 0.5 using a 1/1000 dilution from an overnight preculture. Cells were immobilized in same medium supplemented with 1% agarose and imaged using 100 ms ph and 100 ms GFP acquisition time. Microscopy data reported in this paper will be shared by the lead contact upon request. At submission Microscopy images are available at Mendeley Data at https://data.mendeley.com/preview/rg7db6xyh7?a=31ff605f-f345-4a8e-96b6-b58fa94906d2

### Microscopy Image analysis

Analyses were made with Python version 3.9.18. Cells were automatically segmented in ph images, using Omnipose version 1.0.6 and the available pre-trained model bact_phase_omni. Background was estimated as mean fluorescence of inverted segmentation masks and was subtracted from images. Then cells were segmented automatically using the same Omnipose model as for the quantification of the fluorescence over time. Background was estimated as the mean fluorescence intensity of the inverted cell segmentation masks and was subtracted. A median filter of radius 2 was then applied. Foci were detected with a Laplacian of Gaussian method computed with the scikit-image (version 0.24) method skimage.feature.blob_log() with the following parameter values: min_sigma=1, max_sigma=2, threshold_rel = 0.1. To remove false positive spot detection a filtering step based on estimates of the contrast and the signal to noise ratio (SNR) values of the detected spots was applied. Threshold values were chosen based on the distribution of the spots contrast and SNR values obtained to filter out false spot detections. For time lapse experiment with *E. coli* IbpA-msfGFP, a total of 968 and 262 cells were examined at T=0 and 1256 and 330 cells at T=90, in presence or absence of DMF, respectively. For time lapse experiment with *E. coli* harboring *PahpC*-mut3GFP in presence or absence of DMF, a total of 373 cells and 607 cells were examined at T=0 min, and 458 and 730 cells at T=180 min, for no DMF and DMF conditions, respectively. For time lapse experiment with *E. coli* harboring *PmarR*-mut3GFP in presence or absence of DMF, a total of 87 cells and 267 cells were examined at T=0 min, and 151 and 1008 cells at T=180 min , for no DMF and DMF conditions, respectively.

### Software Data Analysis

Data was analyzed using software described in Reagents or source table.

### pLOGO sequence analysis

We used pLOGO statistical tool to calculate frequency of residues around the Cys^suc^. This tool utilizes a Power Weight Matrix based on binomial probabilities of residue frequencies with respect to proteomic background^18^. Before analysis sequence residues containing Cys^suc^ identified by mass spectrometry as MSC or DSC were separated in one total or two different lists. Sequences were trimmed to produce a nine amino acid sequence including the Cys^suc^, four residues upstream and downstream from the Cys^suc^. This approach generated 140 MSC and 311 DSC individual sequences that were analyzed using pLOGO with the following parameters: remove duplicated sequences, background: *E. coli* K-12, C at -1 fixed. This analysis produced 135 and 300 unique sequences MSC or DSC, respectively.

## QUANTIFICATION AND STATISTICAL ANALYSIS

Sample sizes, reproducibility, and statistical tests used for each figure are denoted in the figure legends or in the method details section. Unless otherwise noted, data presented represent n = 3 independent biological replicates performed. Mass spectrometry data represent n = 5 biological replicates within a single mass spectrometry experiment for DMF-exposed cells or DMSO control. Graphs and images were generated using software and algorithms described in Reagents and resource table.

## ADDITIONAL RESOURCES

No additional resources were used in this study

## REFERENCES

1. Ramazi, S., and Zahiri, J. (2021). Post-translational modifications in proteins: resources, tools and prediction methods. Database: The Journal of Biological Databases and Curation 2021, baab012. 10.1093/database/baab012.

2. Ruecker, N., Jansen, R., Trujillo, C., Puckett, S., Jayachandran, P., Piroli, G.G., Frizzell, N., Molina, H., Rhee, K.Y., and Ehrt, S. (2017). Fumarase Deficiency Causes Protein and Metabolite Succination and Intoxicates Mycobacterium tuberculosis. Cell Chemical Biology 24, 306–315. 10.1016/j.chembiol.2017.01.005.

3. Tomlinson, I.P.M., Alam, N.A., Rowan, A.J., Barclay, E., Jaeger, E.E.M., Kelsell, D., Leigh, I., Gorman, P., Lamlum, H., Rahman, S., et al. (2002). Germline mutations in FH predispose to dominantly inherited uterine fibroids, skin leiomyomata and papillary renal cell cancer. Nat Genet 30, 406–410. 10.1038/ng849.

4. Bresciani, G., Manai, F., Davinelli, S., Tucci, P., Saso, L., and Amadio, M. (2023). Novel potential pharmacological applications of dimethyl fumarate—an overview and update. Front Pharmacol 14, 1264842. 10.3389/fphar.2023.1264842.

5. Werdenberg, D., Joshi, R., Wolffram, S., Merkle, H.P., and Langguth, P. (2003). Presystemic metabolism and intestinal absorption of antipsoriatic fumaric acid esters. Biopharm Drug Dispos 24, 259–273. 10.1002/bdd.364.

6. Kastrati, I., Siklos, M.I., Calderon-Gierszal, E.L., El-Shennawy, L., Georgieva, G., Thayer, E.N., Thatcher, G.R.J., and Frasor, J. (2016). Dimethyl Fumarate Inhibits the Nuclear Factor κB Pathway in Breast Cancer Cells by Covalent Modification of p65 Protein. J Biol Chem 291, 3639–3647. 10.1074/jbc.M115.679704.

7. Kornberg, M.D., Bhargava, P., Kim, P.M., Putluri, V., Snowman, A.M., Putluri, N., Calabresi, P.A., and Snyder, S.H. (2018). Dimethyl fumarate targets GAPDH and aerobic glycolysis to modulate immunity. Science 360, 449–453. 10.1126/science.aan4665.

8. Humphries, F., Shmuel-Galia, L., Ketelut-Carneiro, N., Li, S., Wang, B., Nemmara, V.V., Wilson, R., Jiang, Z., Khalighinejad, F., Muneeruddin, K., et al. (2020). Succination inactivates gasdermin D and blocks pyroptosis. Science 369, 1633–1637. 10.1126/science.abb9818.

9. Schmidt, T.J., Ak, M., and Mrowietz, U. (2007). Reactivity of dimethyl fumarate and methylhydrogen fumarate towards glutathione and N-acetyl-l-cysteine—Preparation of S-substituted thiosuccinic acid esters. Bioorganic & Medicinal Chemistry 15, 333–342. 10.1016/j.bmc.2006.09.053.

10. Bardella, C., El-Bahrawy, M., Frizzell, N., Adam, J., Ternette, N., Hatipoglu, E., Howarth, K., O’Flaherty, L., Roberts, I., Turner, G., et al. (2011). Aberrant succination of proteins in fumarate hydratase-deficient mice and HLRCC patients is a robust biomarker of mutation status. J Pathol 225, 4–11. 10.1002/path.2932.

11. Guberovic, I., and Frezza, C. (2024). Functional implications of fumarate-induced cysteine succination. Trends in Biochemical Sciences, S0968000424001130. 10.1016/j.tibs.2024.05.003.

12. Ferri, C., Castellazzi, M., Merli, N., Laudisi, M., Baldin, E., Baldi, E., Mancabelli, L., Ventura, M., and Pugliatti, M. (2023). Gut Microbiota Changes during Dimethyl Fumarate Treatment in Patients with Multiple Sclerosis. International Journal of Molecular Sciences 24, 2720. 10.3390/ijms24032720.

13. Manai, F., Zanoletti, L., Arfini, D., Micco, S.G.D., Gjyzeli, A., Comincini, S., and Amadio, M. (2023). Dimethyl Fumarate and Intestine: From Main Suspect to Potential Ally against Gut Disorders. International Journal of Molecular Sciences 24, 9912. 10.3390/ijms24129912.

14. Zhou, X., Baumann, R., Gao, X., Mendoza, M., Singh, S., Katz Sand, I., Xia, Z., Cox, L.M., Chitnis, T., Yoon, H., et al. (2022). Gut microbiome of multiple sclerosis patients and paired household healthy controls reveal associations with disease risk and course. Cell 185, 3467–3486.e16. 10.1016/j.cell.2022.08.021.

15. Seta, F.D., Boschi-Muller, S., Vignais, M.L., and Branlant, G. (1997). Characterization of Escherichia coli strains with gapA and gapB genes deleted. Journal of Bacteriology 179, 5218–5221. 10.1128/jb.179.16.5218-5221.1997.

16. Blewett, M.M., Xie, J., Zaro, B.W., Backus, K.M., Altman, A., Teijaro, J.R., and Cravatt, B.F. (2016). Chemical proteomic map of dimethyl fumarate–sensitive cysteines in primary human T cells. Science Signaling 9, rs10–rs10. 10.1126/scisignal.aaf7694.

17. Kulkarni, R.A., Bak, D.W., Wei, D., Bergholtz, S.E., Briney, C.A., Shrimp, J.H., Alpsoy, A., Thorpe, A.L., Bavari, A.E., Crooks, D.R., et al. (2019). A chemoproteomic portrait of the oncometabolite fumarate. Nat Chem Biol 15, 391–400. 10.1038/s41589-018-0217-y.

18. O’Shea, J.P., Chou, M.F., Quader, S.A., Ryan, J.K., Church, G.M., and Schwartz, D. (2013). pLogo: a probabilistic approach to visualizing sequence motifs. Nat Methods 10, 1211–1212. 10.1038/nmeth.2646.

19. Govers, S.K., Mortier, J., Adam, A., and Aertsen, A. (2018). Protein aggregates encode epigenetic memory of stressful encounters in individual Escherichia coli cells. PLoS Biol 16, e2003853. 10.1371/journal.pbio.2003853.

20. Laskowska, E., Wawrzynów, A., and Taylor, A. (1996). IbpA and IbpB, the new heat-shock proteins, bind to endogenous Escherichia coli proteins aggregated intracellularly by heat shock. Biochimie 78, 117–122. 10.1016/0300-9084(96)82643-5.

21. Chiang, S.M., and Schellhorn, H.E. (2012). Regulators of oxidative stress response genes in Escherichia coli and their functional conservation in bacteria. Arch Biochem Biophys 525, 161–169. 10.1016/j.abb.2012.02.007.

22. van der Ploeg, J.R., Weiss, M.A., Saller, E., Nashimoto, H., Saito, N., Kertesz, M.A., and Leisinger, T. (1996). Identification of sulfate starvation-regulated genes in Escherichia coli: a gene cluster involved in the utilization of taurine as a sulfur source. Journal of Bacteriology 178, 5438–5446. 10.1128/jb.178.18.5438-5446.1996.

23. Brugiroux, S., Beutler, M., Pfann, C., Garzetti, D., Ruscheweyh, H.-J., Ring, D., Diehl, M., Herp, S., Lötscher, Y., Hussain, S., et al. (2016). Genome-guided design of a defined mouse microbiota that confers colonization resistance against Salmonella enterica serovar Typhimurium. Nat Microbiol 2, 1–12. 10.1038/nmicrobiol.2016.215.

24. Weiss, A.S., Burrichter, A.G., Durai Raj, A.C., von Strempel, A., Meng, C., Kleigrewe, K., Münch, P.C., Rössler, L., Huber, C., Eisenreich, W., et al. (2022). In vitro interaction network of a synthetic gut bacterial community. ISME J 16, 1095–1109. 10.1038/s41396-021-01153-z.

25. Jones, S.A., Gibson, T., Maltby, R.C., Chowdhury, F.Z., Stewart, V., Cohen, P.S., and Conway, T. (2011). Anaerobic respiration of Escherichia coli in the mouse intestine. Infect. Immun. 79, 4218–4226. 10.1128/IAI.05395-11.

26. Manuel, A.M., Walla, M.D., Faccenda, A., Martin, S.L., Tanis, R.M., Piroli, G.G., Adam, J., Kantor, B., Mutus, B., Townsend, D.M., et al. (2017). Succination of Protein Disulfide Isomerase Links Mitochondrial Stress and Endoplasmic Reticulum Stress in the Adipocyte During Diabetes. Antioxidants & Redox Signaling 27, 1281–1296. 10.1089/ars.2016.6853.

27. Kamariah, N., Sek, M.F., Eisenhaber, B., Eisenhaber, F., and Grüber, G. (2016). Transition steps in peroxide reduction and a molecular switch for peroxide robustness of prokaryotic peroxiredoxins. Scientific Reports 6, 37610. 10.1038/srep37610.

28. Yun, M., Park, C.-G., Kim, J.-Y., and Park, H.-W. (2000). Structural Analysis of Glyceraldehyde 3-Phosphate Dehydrogenase from Escherichia coli: Direct Evidence of Substrate Binding and Cofactor-Induced Conformational Changes,. Biochemistry 39, 10702–10710. 10.1021/bi9927080.

29. Niehaus, T.D., Folz, J., McCarty, D.R., Cooper, A.J.L., Moraga Amador, D., Fiehn, O., and Hanson, A.D. (2018). Identification of a metabolic disposal route for the oncometabolite S-(2-succino)cysteine in Bacillus subtilis. J Biol Chem 293, 8255–8263. 10.1074/jbc.RA118.002925.

30. Hillmann, K.B., Goethel, M.E., Erickson, N.A., and Niehaus, T.D. (2022). Identification of a S-(2-succino)cysteine breakdown pathway that uses a novel S-(2-succino) lyase. J Biol Chem 298, 102639. 10.1016/j.jbc.2022.102639.

31. Matthews, A., Schönfelder, J., Lagies, S., Schleicher, E., Kammerer, B., Ellis, H.R., Stull, F., and Teufel, R. (2022). Bacterial flavoprotein monooxygenase YxeK salvages toxic S-(2-succino)-adducts via oxygenolytic C–S bond cleavage. The FEBS Journal 289, 787–807. 10.1111/febs.16193.

32. Eberl, C., Weiss, A.S., Jochum, L.M., Raj, A.C.D., Ring, D., Hussain, S., Herp, S., Meng, C., Kleigrewe, K., Gigl, M., et al. (2021). E. coli enhance colonization resistance against Salmonella Typhimurium by competing for galactitol, a context-dependent limiting carbon source. Cell Host & Microbe 29, 1680–1692.e7. 10.1016/j.chom.2021.09.004.

33. Lénon, M., Arias-Cartín, R., and Barras, F. (2022). The Fe–S proteome of Escherichia coli: prediction, function, and fate. Metallomics 14, mfac022. 10.1093/mtomcs/mfac022.

34. Py, B., and Barras, F. (2010). Building Fe–S proteins: bacterial strategies. Nat Rev Microbiol 8, 436–446. 10.1038/nrmicro2356.

35. Bachmann, B.J. (1987). Derivations and genotypes of some mutant derivatives of Escherichia coli K12. In.

36. Lagkouvardos, I., Pukall, R., Abt, B., Foesel, B.U., Meier-Kolthoff, J.P., Kumar, N., Bresciani, A., Martínez, I., Just, S., Ziegler, C., et al. (2016). The Mouse Intestinal Bacterial Collection (miBC) provides host-specific insight into cultured diversity and functional potential of the gut microbiota. Nat Microbiol 1, 1–15. 10.1038/nmicrobiol.2016.131.

37. Auchtung, J.M., Robinson, C.D., Farrell, K., and Britton, R.A. (2016). MiniBioReactor Arrays (MBRAs) as a Tool for Studying C. difficile Physiology in the Presence of a Complex Community. In Clostridium difficile: Methods and Protocols, A. P. Roberts and P. Mullany, eds. (Springer), pp. 235–258. 10.1007/978-1-4939-6361-4_18.

38. Elbing, K., and Brent, R. (2019). Recipes and tools for culture of Escherichia coli. Curr Protoc Mol Biol 125, e83. 10.1002/cpmb.83.

39. Zaslaver, A., Bren, A., Ronen, M., Itzkovitz, S., Kikoin, I., Shavit, S., Liebermeister, W., Surette, M.G., and Alon, U. (2006). A comprehensive library of fluorescent transcriptional reporters for Escherichia coli. Nat Methods 3, 623–628. 10.1038/nmeth895.

40. Deghelt, M., Cho, S.-H., Govers, S.K., Janssens, A., Dachsbeck, A., Remaut, H.K., and Collet, J.-F. (2023). The outer membrane and peptidoglycan layer form a single mechanical device balancing turgor (Microbiology) 10.1101/2023.04.29.538579.

41. Wahl, A., Hubert, P., Sturgis, J.N., and Bouveret, E. (2009). Tagging of Escherichia coli proteins with new cassettes allowing in vivo systematic fluorescent and luminescent detection, and purification from physiological expression levels. PROTEOMICS 9, 5389–5393. 10.1002/pmic.200900240.

42. Wahl, A., My, L., Dumoulin, R., Sturgis, J.N., and Bouveret, E. (2011). Antagonistic regulation of dgkA and plsB genes of phospholipid synthesis by multiple stress responses in Escherichia coli. Molecular Microbiology 80, 1260–1275. 10.1111/j.1365-2958.2011.07641.x.

43. Loiseau, L., Ollagnier-de Choudens, S., Lascoux, D., Forest, E., Fontecave, M., and Barras, F. (2005). Analysis of the heteromeric CsdA-CsdE cysteine desulfurase, assisting Fe-S cluster biogenesis in Escherichia coli. J. Biol. Chem. 280, 26760–26769. 10.1074/jbc.M504067200.

44. Datsenko, K.A., and Wanner, B.L. (2000). One-step inactivation of chromosomal genes in Escherichia coli K-12 using PCR products. Proceedings of the National Academy of Sciences 97, 6640–6645. 10.1073/pnas.120163297.

45. Schindelin, J., Arganda-Carreras, I., Frise, E., Kaynig, V., Longair, M., Pietzsch, T., Preibisch, S., Rueden, C., Saalfeld, S., Schmid, B., et al. (2012). Fiji: an open-source platform for biological-image analysis. Nat Methods *9*, 676–682. 10.1038/nmeth.2019.

46. Cutler, K.J., Stringer, C., Lo, T.W., Rappez, L., Stroustrup, N., Brook Peterson, S., Wiggins, P.A., and Mougous, J.D. (2022). Omnipose: a high-precision morphology-independent solution for bacterial cell segmentation. Nat Methods 19, 1438–1448. 10.1038/s41592-022-01639-4.

47. Szklarczyk, D., Kirsch, R., Koutrouli, M., Nastou, K., Mehryary, F., Hachilif, R., Gable, A.L., Fang, T., Doncheva, N.T., Pyysalo, S., et al. (2023). The STRING database in 2023: protein-protein association networks and functional enrichment analyses for any sequenced genome of interest. Nucleic Acids Res 51, D638–D646. 10.1093/nar/gkac1000.

48. Tyanova, S., Temu, T., and Cox, J. (2016). The MaxQuant computational platform for mass spectrometry-based shotgun proteomics. Nat Protoc 11, 2301–2319. 10.1038/nprot.2016.136.

49. Cox, J., Neuhauser, N., Michalski, A., Scheltema, R.A., Olsen, J.V., and Mann, M. (2011). Andromeda: a peptide search engine integrated into the MaxQuant environment. J Proteome Res 10, 1794–1805. 10.1021/pr101065j.

50. The UniProt Consortium (2025). UniProt: the Universal Protein Knowledgebase in 2025. Nucleic Acids Research 53, D609–D617. 10.1093/nar/gkae1010.

51. Bolyen, E., Rideout, J.R., Dillon, M.R., Bokulich, N.A., Abnet, C.C., Al-Ghalith, G.A., Alexander, H., Alm, E.J., Arumugam, M., Asnicar, F., et al. (2019). Reproducible, interactive, scalable and extensible microbiome data science using QIIME 2. Nat Biotechnol 37, 852–857. 10.1038/s41587-019-0209-9.

52. Turnbaugh, P.J., Hamady, M., Yatsunenko, T., Cantarel, B.L., Duncan, A., Ley, R.E., Sogin, M.L., Jones, W.J., Roe, B.A., Affourtit, J.P., et al. (2009). A core gut microbiome in obese and lean twins. Nature 457, 480–484. 10.1038/nature07540.

53. Auchtung, J.M., Robinson, C.D., Farrell, K., and Britton, R.A. (2016). MiniBioReactor Arrays (MBRAs) as a Tool for Studying C. difficile Physiology in the Presence of a Complex Community. In Clostridium difficile Methods in Molecular Biology., A. P. Roberts and P. Mullany, eds. (Springer New York), pp. 235–258. 10.1007/978-1-4939-6361-4_18.

54. Thompson, L.R., Sanders, J.G., McDonald, D., Amir, A., Ladau, J., Locey, K.J., Prill, R.J., Tripathi, A., Gibbons, S.M., Ackermann, G., et al. (2017). A communal catalogue reveals Earth’s multiscale microbial diversity. Nature 551, 457–463. 10.1038/nature24621.

55. Osborn, M.J., Gander, J.E., Parisi, E., and Carson, J. (1972). Mechanism of assembly of the outer membrane of Salmonella typhimurium. Isolation and characterization of cytoplasmic and outer membrane. J Biol Chem 247, 3962–3972.

56. Hausladen, A., and Fridovich, I. (1996). Measuring nitric oxide and superoxide: rate constants for aconitase reactivity. Methods Enzymol 269, 37–41. 10.1016/s0076-6879(96)69007-7.

57. Gort, A.S., and Imlay, J.A. (1998). Balance between Endogenous Superoxide Stress and Antioxidant Defenses. Journal of Bacteriology 180, 1402–1410.

58. Jones, A.J.Y., and Hirst, J. (2013). A spectrophotometric coupled enzyme assay to measure the activity of succinate dehydrogenase. Analytical Biochemistry 442, 19–23. 10.1016/j.ab.2013.07.018.

59. Hughes, C.S., Foehr, S., Garfield, D.A., Furlong, E.E., Steinmetz, L.M., and Krijgsveld, J. (2014). Ultrasensitive proteome analysis using paramagnetic bead technology. Mol Syst Biol 10, 757. 10.15252/msb.20145625.

60. Merkley, E.D., Metz, T.O., Smith, R.D., Baynes, J.W., and Frizzell, N. (2014). The succinated proteome. Mass Spectrom Rev 33, 98–109. 10.1002/mas.21382.

61. Cox, J., Hein, M.Y., Luber, C.A., Paron, I., Nagaraj, N., and Mann, M. (2014). Accurate Proteome-wide Label-free Quantification by Delayed Normalization and Maximal Peptide Ratio Extraction, Termed MaxLFQ. Mol Cell Proteomics 13, 2513–2526. 10.1074/mcp.M113.031591.

62. Perez-Riverol, Y., Bandla, C., Kundu, D.J., Kamatchinathan, S., Bai, J., Hewapathirana, S., John, N.S., Prakash, A., Walzer, M., Wang, S., et al. (2025). The PRIDE database at 20 years: 2025 update. Nucleic Acids Res 53, D543–D553. 10.1093/nar/gkae1011.

63. Ritchie, M.E., Phipson, B., Wu, D., Hu, Y., Law, C.W., Shi, W., and Smyth, G.K. (2015). limma powers differential expression analyses for RNA-sequencing and microarray studies. Nucleic Acids Res 43, e47. 10.1093/nar/gkv007.

64. Giai Gianetto, Q., Combes, F., Ramus, C., Bruley, C., Couté, Y., and Burger, T. (2016). Calibration plot for proteomics: A graphical tool to visually check the assumptions underlying FDR control in quantitative experiments. Proteomics 16, 29–32. 10.1002/pmic.201500189.

